# Random guide-independent DNA cleavage from the Argonaute of *Exiguobacterium* sp. AB2

**DOI:** 10.1101/2025.02.21.639407

**Authors:** Miguel Antonio M. Cañiza, Ron Leonard V. Dy

## Abstract

**Background:** Bacteria and bacteriophages (phages) are locked in a coevolutionary “arms race” to outcompete one another with novel systems and strategies. Regularly outnumbered tenfold by phages, bacteria have responded to the constant threat of phage predation by evolving a vast array of sophisticated defense systems. Among these, prokaryotic Argonautes (pAgos) are nucleic acid-guided endonucleases that target complementary sequences of invading mobile genetic elements (MGEs). However, as the preference for targeting MGE sequences has been demonstrated in only a limited number of pAgos, their precise physiological functions remain elusive. Here, we discovered a pAgo in *Exiguobacterium* sp. AB2, EsAgo, encoded in close proximity to other putative defense systems on the *E*. AB2 genome. Such clustering into genomic “defense islands” is a common phenomenon among prokaryotic defense systems, further implicating pAgos with a role in host defense. Accordingly, we had sought to characterize EsAgo as a nucleic acid-guided nucleic acid-targeting nuclease against MGEs for bacterial defense in this study.

**Results:** Using sequence to structure homology tools, we show that the predicted model of EsAgo exhibits the structural characteristics typical of a full-length, catalytically active, DNA-guided pAgo. Akin to other pAgos, EsAgo uses a divalent cation cofactor to indiscriminately “chop” plasmids *in vitro*. Furthermore, a site-directed double mutant of EsAgo bearing two missense mutations at the catalytic site exhibited significantly reduced levels of this random plasmid-degrading activity. Lastly, when EsAgo was supplied with synthetic 5’-P ssDNA guides, random nuclease activity was attenuated and may have resulted in the sequence-specific cleavage of dsDNA at 37°C.

**Conclusions:** These findings suggest that EsAgo functions as a DNA-interfering nuclease with or without DNA guides. Within the cell, it is possible that EsAgo utilizes this mechanism to screen and destroy foreign genetic elements. Moreover, the potential capacity for specific dsDNA cleavage at moderate temperatures gives rise to intriguing possibilities of repurposing EsAgo as a programmable nuclease for future biotechnological use.

## BACKGROUND

Argonaute proteins (Agos) are present in all domains of life [1, 2, 3]. Fundamentally, Agos are characterized by their ability to target a specific nucleic acid sequence using a short (∼15-30 nt) and complementary nucleic acid guide strand [2]. This capability for sequence-specific recognition is sometimes coupled with the cleavage of the complementary target by the Ago itself. These proteins were first discovered in eukaryotes [4] where they play a central role in RNA interference (RNAi) pathways as the catalytic core of the RNA-induced silencing complex (RISC) [5, 6, 7]. Accordingly, Agos are encoded in the majority (∼65%) of sequenced eukaryotic genomes [1]. Interestingly, prokaryotic homologs of eukaryotic Agos (eAgos) are also present in ∼32% of sequenced archaeal genomes and ∼9% of sequenced bacterial genomes [1].

Phylogenetic analyses of Agos suggest their origin in prokaryotes [1]. It is believed that pAgos were inherited among various prokaryotes through extensive horizontal gene transfer (HGT), whereas eAgos appear to have evolved solely through vertical inheritance [1, 8, 9]. Consequently, pAgos are highly diverse proteins and form three phylogenetic clades based on their domain organization: long-A, long-B, and short pAgos [2, 10]. Long (i.e. full-length) pAgos possess all four domains found in eAgos: N (amino-terminal), PAZ (<u>P</u>IWI-<u>A</u>rgonaute-<u>Z</u>wille), MID (middle), and PIWI (<u>P</u>-element Induced <u>Wi</u>mpy Testis). On the other hand, short pAgos typically possess a MID and PIWI [2]. The N and PAZ are less conserved among pAgos with the former domain believed to facilitate guide-target duplex unwinding while the latter is responsible for binding the 3’ end of the guide strand [1, 2, 10]. The MID contains a binding pocket for the 5’ end of the guide comprised of a divalent cation and a conserved amino acid motif, predominantly **YKQ**TN**K** (i.e. YK-type, most conserved residues in bold) among pAgos that preferentially utilize 5’-phosphorylated (5’-P) guides [2]. Nuclease activity is derived from a catalytic tetrad of highly conserved residues in the PIWI domain: DEDX where X represents an aspartate, histidine, or a lysine [1, 2, 10, 11]. Generally, long-A pAgos contain complete DEDX tetrads, whereas long-B and short pAgos exclusively possess catalytically inactive PIWI domains [2].

The similarities between eAgo and long pAgo domain constitution suggests that their activity as guided nucleases is maintained despite divergent evolutionary histories [1]. However, as prokaryotes lack RNAi pathways homologous to those found in eukaryotes [12], this suggests that the physiological function of pAgos is distinct from that of their eukaryotic counterparts. Instead of regulating mRNA transcript levels, targeted interference against plasmid and bacteriophage DNA has been broadly observed in a number of pAgos—the majority of which belong to the long-A clade [3]. For example, *Thermus thermophilus* Ago (TtAgo) [13], *Pyrococcus furiosus* Ago (PfAgo) [14], and *Rhodobacter sphaeroides* Ago (RsAgo) [15] can facilitate the targeted degradation of plasmid DNA within their native bacterial hosts. Furthermore, some pAgos expressed in heterologous hosts such as *Methanocaldococcus janaschii* Ago (MjAgo) reduce plasmid transformation efficiency [16] or otherwise preferentially acquire plasmid-derived ssDNA guides such as CbAgo from *Clostridium butyricum* Ago (CbAgo) [17] and *Exiguobacterium marinum* Ago (EmaAgo) [18], among others [18]. Direct interference against phages is less documented with only two reported cases: CbAgo [19] and EmaAgo [18] protecting *Escherichia coli* from M13 and P1vir coliphage infection, respectively. In contrast, the catalytically inactive short pAgos provide population-based immunity by functioning as abortive infection (Abi) systems which activate associated proteins such as nonspecific nucleases [20], NAD(P)ases [21], and depolarization-inducing membrane enzymes [22] upon guide-mediated recognition of foreign genetic elements [3]. While such evidence implies that pAgos confer immunity to MGE infection, more experimental validation is needed to confirm this as these studies are among only a handful detailing pAgo-mediated MGE interference *in vivo*.

As postulated components of the prokaryotic immune system, this raises the question: how do pAgos discriminate between self (i.e. genome) and non-self genetic elements? At present this remains poorly understood as no dedicated pAgo guide biogenesis pathway has been identified [3]. One hypothesis is that multicopy sequences undergoing replication may be recognized as “foreign” and are selectively processed by pAgos for functional guides due to the greater availability of substrate DNA [15]. In such cases, it is assumed that long-A pAgos generate their own guides as no pre-processing enzymes analogous to Dicer and Drosha have been identified in prokaryotes [16]. Here, the apoenzyme form of some catalytically active long-A pAgos such as TtAgo [23], CbAgo [24], *Limnothrix rosea* Ago (LrAgo) [24], and MjAgo [16] are capable of degrading plasmid dsDNA in a guide-independent “chopping” manner, albeit inefficiently, whereupon smaller degraded ssDNA fragments may be loaded onto the pAgo as guides [3].

Bacteria are not protected from MGEs by pAgos alone. On average, most bacteria possess about five or six distinct antiviral defense systems [25, 26, 27]. Interestingly, the genes encoding these systems tend to cluster together in genomic “defense islands”, within which *restriction-modification* (RM), *toxin-antitoxin* (TA), *Abi*, and *pAgo* systems are commonly found [10, 28, 29, 30]. It is unclear why such clustering occurs; however, the characterization of unknown genes residing in these “defense islands” has led to the discovery of numerous antiphage defenses [25, 30, 31].

In 2014, Cabria *et al*. [32] isolated a novel mesophilic *Exiguobacterium* species, AB2, from Manleluag ophiolitic spring, Pangasinan, Philippines. Following the sequencing, assembly, and annotation of its genome [32], we identified a putative defense island containing known defense systems such a type II TA, type II Wadjet, type I RM, and a <u>d</u>efense island <u>s</u>ystem <u>a</u>ssociated with restriction-<u>m</u>odification (DISARM) system (**Fig. 1**). Interestingly, we identified a novel full-length pAgo, *EsAgo* (*<u>E</u>xiguobacterium* <u>sp</u>. AB2 <u>Ago</u>), about ∼10.8 kb downstream of the DISARM system. We had therefore hypothesized that EsAgo functions for host defense against MGE infection and combine *in silico*, *in vivo*, and *in vitro* characterization of this protein to probe its capacity for nucleic acid-guided nucleic acid-targeting in this study.

**Fig. 1.**
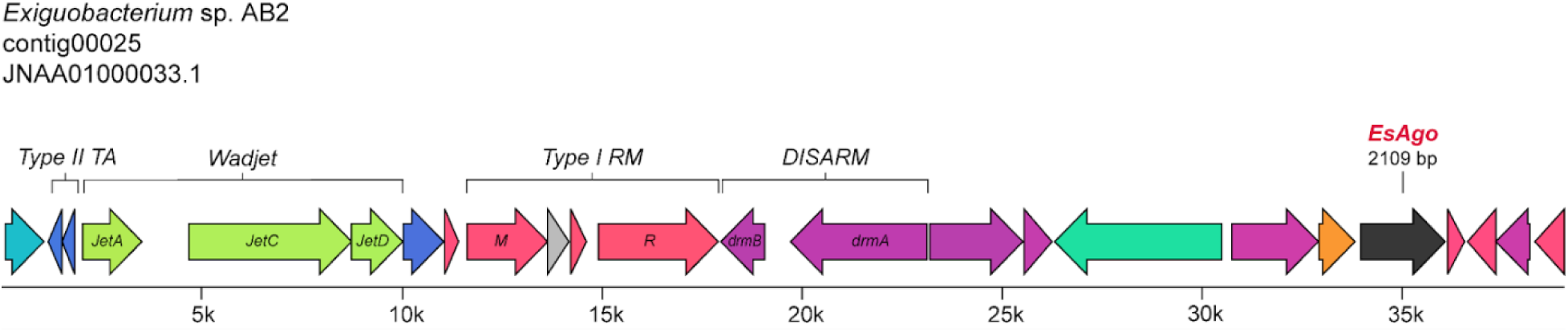
Genomic neighborhood of EsAgo. *EsAgo* is located on contig00025 of the *E*. AB2 genome which can be accessed at NCBI GenBank accession no. JNA01000033.1. Annotation of this contig revealed the presence of coding sequences (CDS) of known genes and those with domains of unknown function (DUF). Known defense systems include a type II TA, a Wadjet (composed of the genes *jetACD*), a type I RM (composed of separate <u>M</u>ethylation and <u>R</u>estriction subunits), and a DISARM system (composed of the genes *drmAB*). *EsAgo* (accession no. WP_034806158.1) is located further downstream of these genes at position (+)33,959-36,067 of the contig.

## METHODS

### Multiple genome alignment and annotation of *Exiguobacterium* contigs

To further validate the placement of *EsAgo* in a genomic “defense island” on the *E.* AB2 genome, the immediate genomic neighborhoods of *EsAgo* and other *pAgos* derived from various *Exiguobacterium* species were compared. To identify other *Exiguobacterium* pAgos, a PSI-BLAST of the protein sequence of EsAgo was first performed. Here, the individual contigs containing *EsAgo* and other *Exiguobacterium* pAgo genes were aligned against each other using clinker (default parameters) [33]. To identify known defense systems residing on these contigs, these were submitted to the <u>P</u>rokaryotic <u>A</u>ntiviral <u>D</u>efense <u>LOC</u>ator (PADLOC) web server (https://padloc.otago.ac.nz/padloc/, Accessed: September 2023, April 2024) [34]. The detection of non-coding RNAs, such as CRISPR arrays, was excluded from this analysis as this substantially increased runtime [34]. It should also be noted that TA systems are not included in the PADLOC database [34]. The TA system of *E.* AB2 was identified via BLASTX/P.

### Multiple sequence alignment of pAgos

Protein sequences of pAgos from other *Exiguobacterium* species were aligned against that of EsAgo using the Geneious aligner algorithm tool integrated into Geneious Prime® v2023.0.4. Protein sequences of experimentally characterized pAgos from various genera were similarly aligned against EsAgo wherein the overwhelming majority of these were of the long-A type described by Ryazansky *et al*. [2]. The corresponding phylogenetic trees were also generated by the Geneious aligner algorithm integrated into Geneious Prime® v2023.0.4 from these multiple sequence alignments.

### Protein modeling

The protein sequence of EsAgo was submitted to the PHYRE2 Protein Fold Recognition Server (https://www.sbg.bio.ic.ac.uk/~phyre2/html/page.cgi?id=index, Accessed: November 2, 2020) to generate a homology detection-based three-dimensional model under “intensive” conditions [35]. *De novo* protein structure prediction (i.e. no homologous structural templates) was also performed using the simplified AlphaFold v2.3.2 algorithm on AlphaFold Colab (Colab notebook) (https://colab.research.google.com/github/deepmind/alphafold/blob/main/notebooks/AlphaFold.ipynb, Accessed: July 24, 2022) [36]. Default parameters were used with the exception of disabling the relaxation stage (run would not complete with this enabled) which may have consequently introduced distracting small stereochemical violations into the AlphaFold model [36]. Generated protein models were viewed and compared using PyMOL v2.5.3 [37]. Crystal structures of other pAgos were taken from the Research Collaboratory for Structural Bioinformatics Protein Data Bank (RCSB PDB) (https://www.rcsb.org/?ref=nav_home, Accessed: September 09, 2020) and compared to the EsAgo models.

### Bacterial strains and culture conditions

Bacterial cultures of *E*. AB2 were grown at 37°C in Tryptic Soy Broth (TSB) pH 10 at 225 rpm or on solid 1.5% (w/v) Tryptic Soy Agar (TSA) plates. Bacterial cultures of *E*. *coli* were grown at 37°C in Luria-Bertani (LB) broth with 225 rpm shaking or on solid 1.5% (w/v) LB agar (LBA) plates, with antibiotics (100 µg/mL Ap) and/or supplements (1 mM IPTG) when necessary. Optical density at 600 nm (OD_600_) of liquid cultures was measured via a NanoDrop^TM^ 2000 spectrophotometer (Thermo Scientific^TM^). All liquid cultures used in this study were seeded to a starting OD_600_ of 0.05 from a prior overnight culture. Overnight cultures in this study constitute a single bacterial colony inoculated into 5 mL of media with the appropriate antibiotics and supplements and grown for 12-16 hours.

### Protein expression and plasmid shearing assay

To heterologously overexpress EsAgo as no genetic manipulation techniques are available for *E.* AB2, WT *EsAgo* was directly amplified from extracted *E.* AB2 gDNA, cloned into the T5-lacO-based expression vector pQE-80L (generating pSML0001), and transformed into chemically competent *E. coli* DH5ɑ. Additionally, overlap extension PCR was used to introduce aspartic acid to alanine substitutions within the <u>D</u>E<u>D</u>X catalytic tetrad at residues 495 and 564 (i.e. D495A, D564A). This double mutant (DM) *EsAgo* construct was similarly cloned into pQE-80L (generating pSML0050) and introduced into *E. coli* DH5ɑ.

To assess plasmid quality in response to the overexpression of EsAgo WT or DM, *E. coli* cells carrying pSML0001 or pSML0050 were first grown in liquid culture as described above with IPTG added to a working concentration of 1 mM when OD_600_ reached 0.6-0.8 (about 3 hours of growth). Uninduced flasks without IPTG served as negative controls. Proteins were overexpressed for a total of 5 hours at 37°C. Concurrently, plasmids were extracted via miniprep (Zymo) from aliquots at specified hourly intervals (I_0_-I_5_) during IPTG induction. Equal amounts of purified plasmids were resolved on 1.5% (w/v) agarose gels stained with GelGreen® (Biotium) and visualized on an Azure 200® gel doc (Azure Biosystems, catalog no. AZI200-01).

### Purification of His_6_-EsAgo WT and DM

Large volumes of protein were purified by affinity chromatography and size exclusion chromatography (SEC) using an ÄKTA pure^TM^ 25 L1 (Cytiva, catalog no. 29018224) chromatography system. To enable purification, His_6_-tags were added to the N-terminal ends of *EsAgo* WT and DM via in frame cloning into pQE-80L (generating pSML0035 and pSML0053 expression vectors, respectively). Plasmids were transformed into *E. coli* DH5ɑ and cells grown in liquid culture as described above. Protein overexpression was induced with 1 mM IPTG for 5 hours at 37°C when OD_600_ of liquid cultures reached 0.6-0.8. Afterwards, cells were pelleted by centrifugation (2300 x g, 20 minutes, 4°C) and stored overnight at -80°C. Cell pellets were resuspended in Lysis/Binding Buffer (50 mM NaH_2_PO_4_, 300 mM NaCl, 10 mM Imidazole, pH 8.0), supplemented with 1 mg/mL lysozyme and 0.1 mM PMSF, then lysed via 6 sonication cycles (30 seconds on, 30 seconds off, high setting) (Bioruptor Plus® sonicator). Lysate was cleared by centrifugation (10000 x g, 30 minutes, 4°C) with the resulting supernatant (containing soluble proteins) filtered through a 0.22 µm syringe filter to remove dust.

For affinity chromatography, the supernatant was loaded onto a 1 mL HisTrap^TM^HP column (Cytiva) equilibrated with 5 column volumes of Lysis/Binding Buffer. The column was then washed with 15 column volumes of Wash Buffer (50 mM NaH_2_PO_4_, 300 mM NaCl, 40 mM Imidazole, pH 8.0) and His_6_-tagged proteins eluted isocratically with 10 column volumes of Elution Buffer (50 mM NaH_2_PO_4_, 300 mM NaCl, 250 mM Imidazole, pH 8.0). Fractions containing the protein of interest were pooled and concentrated by centrifugation (4000 x g, 1 hour, 4°C) using an Amicon® Ultra Centrifugal Filter (Millipore) with a 3 kDa cut-off. For SEC, concentrated samples were loaded onto a Superdex 200 Increase 10/300 GL column (Cytiva) equilibrated with 1.2 column volumes of SEC Buffer (50 mM NaH_2_PO_4_, 300 mM NaCl, pH 8.0). Proteins of interest were eluted with SEC Buffer, flash frozen in liquid nitrogen or diluted 50% v/v in 100% glycerol, and stored at -80°C. Protein concentrations of eluted fractions were determined using a NanoDrop^TM^ 2000 spectrophotometer (Thermo Scientific^TM^).

### EsAgo DNA cleavage assays

To determine nuclease activity against dsDNA substrates, purified His_6_-tagged EsAgo WT and EsAgo DM were incubated with pQE-80L. As we had tested various reaction conditions, a typical unguided activity reaction consisted of 30 pmols EsAgo mixed with 500 ng pQE-80L (1 pmol EsAgo:∼16.67 ng pQE-80L), unless otherwise indicated. These were incubated at 37°C for 5 hours in 2X Reaction Buffer (working concentration of 10 mM Tris-HCl, 50 mM NaCl, 1 mM Mn^2+^, pH 8.0), unless otherwise indicated. Reaction products were resolved on 1.5% w/v agarose gels stained with GelGreen® (Biotium) and visualized with an Azure 200® gel doc (Azure Biosystems). Gels were analyzed with ImageJ and Graphpad Prism 9.0.0.

For site-specific cleavage of dsDNA at the *lacI* gene of pQE-80L, synthetic 5’-P ssDNA oligonucleotides (oligos) 21 nt in length were supplied at a guide:EsAgo molar ratio of 1:1 and 1:3 for forward and reverse guides, respectively, so as to target opposing DNA strands. In two half reactions, 5 pmols of EsAgo in Reaction Buffer were loaded with either forward or reverse guides via incubation at 37°C for 30 minutes. Subsequently, half reactions were pooled to which 100 ng of pQE-80L was added (1 pmol EsAgo:10 ng pQE-80L) followed by incubation for 5 hours at 37°C. HindIII was added afterwards following the 5-hour incubation period and incubated an additional hour at 37°C to cleave linearized pQE-80L into two fragments of known length. For comparison, this assay was repeated using synthetic 5’-OH ssDNA guides which were otherwise identical in sequence. Reaction products were resolved on 1.5% w/v agarose gels stained with GelGreen® (Biotium) and visualized with an Azure 200® gel doc (Azure Biosystems). Gels were analyzed with ImageJ and Graphpad Prism 9.0.0.

### Statistical Analysis

Statistical analysis was performed using GraphPad Prism version 9.0.0 software with quantified data expressed as means ± standard deviation (SD).

## RESULTS

### Other *Exiguobacterium* pAgos are similarly organized into genomic defense islands

A PSI-BLAST of the protein sequence of EsAgo revealed 25 unique pAgo sequences from

29 *Exiguobacterium* species (some species possess identical pAgos). A phylogenetic tree generated from the multiple protein sequence alignment of EsAgo with these *Exiguobacterium* pAgos is illustrated in **Fig. 2A**. Over half (14 of 25) share a >90% protein sequence identity with EsAgo. The pAgos from *E.* sp. SL-10 and *aestuarii* represent the most closely (97.01% identity) and distantly (87.89% identity) related pAgos, respectively, to EsAgo among those included in this analysis (**Fig. 2A**). Of note as well is that the pAgo from *E. marinum* (EmaAgo), which was recently experimentally characterized by Lisitskaya *et al*. [18], shares an 89.46% sequence identity with EsAgo. Accession numbers of the protein sequences of these *Exiguobacterium* pAgos, as well as their % sequence identity with EsAgo, are listed in **Supplementary Table 1**.

**Fig 2.**
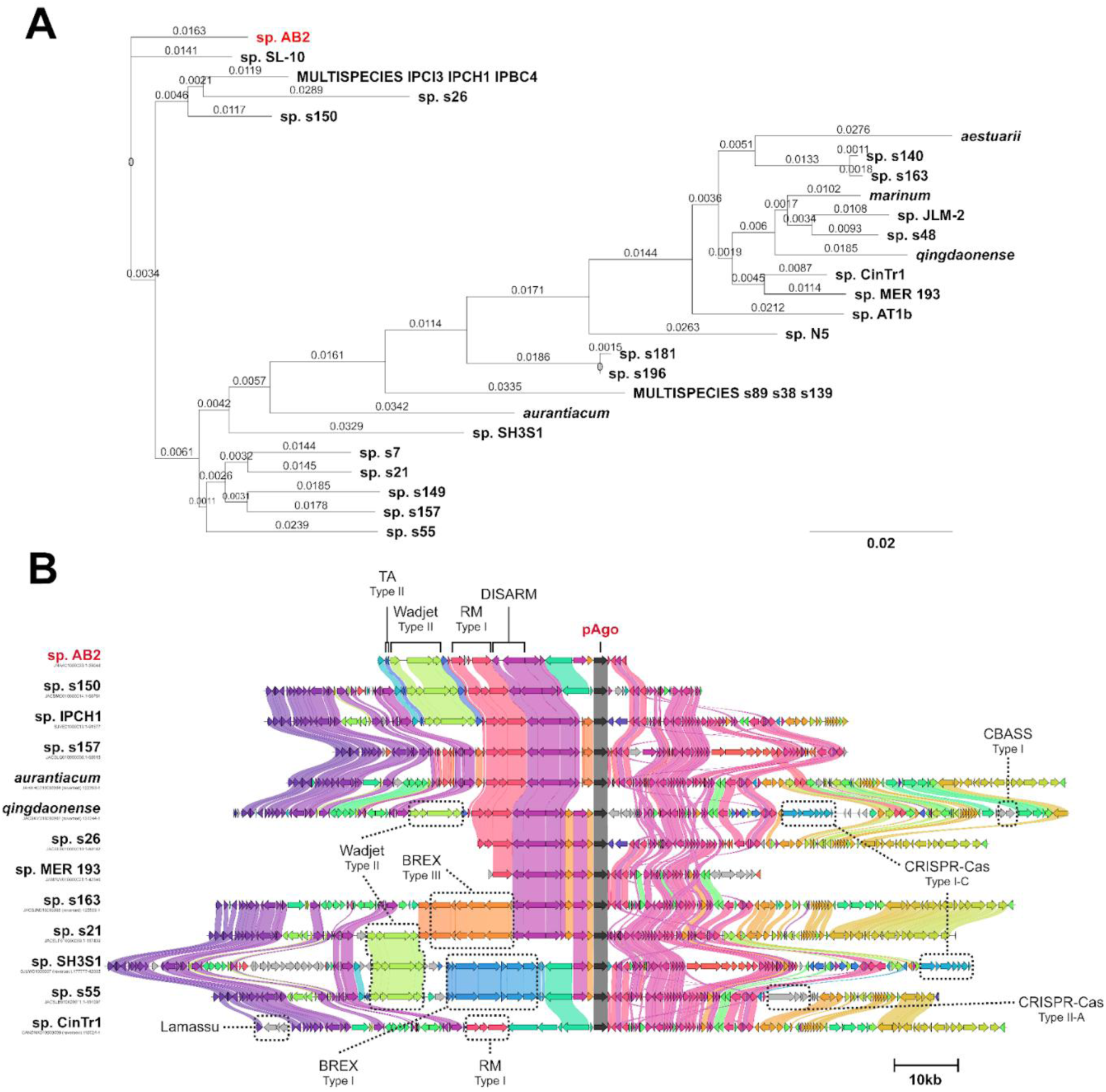
Genomic neighborhoods of *Exiguobacterium* pAgos. (**A**) Neighbor-joining phylogenetic tree (global alignment, BLOSUM80 cost matrix, EsAgo as outgroup) generated from the multiple protein sequence alignment of EsAgo with other *Exiguobacterium* pAgos. Branch labels display substitutions per site. (**B**) Multiple genome alignment of contigs containing *Exiguobacterium* pAgos. The alignment is centered around the pAgo gene (dark gray arrows) while other defense systems present are indicated. NCBI accession numbers of the contigs are displayed under the *Exiguobacterium* species it belongs to and are listed in **Supplementary Table 1**. Contig alignment and gene cluster similarity were generated using clinker [33]. Defense systems were identified using PADLOC [34].

To examine and compare the immediate genomic neighborhoods of these *Exiguobacterium* pAgos, gene cluster alignments were performed on the contigs containing these pAgos using clinker [33]. The contigs which share a similar overall organization to the contig containing *EsAgo* are displayed in **Fig. 2B**. To identify known defense systems residing on these contigs, these were submitted to the PADLOC web server [34]. Similar type II TA, type II Wadjet, type I RM, and/or DISARM systems were annotated on these contigs (**Fig. 2B**). Other defenses identified by PADLOC include a type I <u>c</u>yclic oligonucleotide-<u>b</u>ased <u>a</u>ntiphage <u>s</u>ignaling <u>s</u>ystem (CBASS), type I-C and type II-A CRISPR-Cas systems, type I and type III BREX (Bacteriophage Exclusion) systems, and a Lamassu system (**Fig. 2B**). Interestingly, we had also identified a type I-C CRISPR-Cas system residing on a separate contig on the *E.* AB2 genome (NCBI Accession No. JNAA01000068) (data not shown). Accordingly, it is possible that the region downstream of *EsAgo* may resemble that of *E. qingdaonense* or sp. SH3S1 due to the presence of similar type I-C CRISPR-Cas systems. An expanded view of **Fig. 2B** depicting the multiple genome alignment of all 25 *Exiguobacterium* contigs used in this study, as well as any putative defense systems identified by PADLOC on these contigs, is presented in **Supplementary Fig. 1.** The location of each putative defense system on its respective contig is listed in **Supplementary Table 2**.

### EsAgo exhibits the structural characteristics typical of catalytically active 5’-P DNA-binding long-A pAgos

To determine if *EsAgo* indeed encodes a bona fide pAgo, the protein sequence of EsAgo was aligned with known and characterized pAgos. By extension, this analysis also served to identify any conserved motifs necessary for catalytic activity. The phylogenetic tree generated from this alignment is illustrated in **Fig. 3A** and includes pAgos from different genera. Most pAgos in this alignment were derived from mesophilic organisms with several of these having exhibited DNA-guided cleavage of DNA substrates at moderate temperatures in previous studies. Among these characterized pAgos, the three which share the closest homology to EsAgo are *Clostridium sartagoforme* Ago (CsAgo, 36.3% identity), *Intestinibacter bartlettii* Ago (IbAgo, 32.2% identity), and *Kurthia massiliensis* Ago (KmAgo, 32.0% identity). Additionally, the well-characterized CbAgo and TtAgo share a 21.6% and a 28.4% sequence identity with EsAgo, respectively.

**Fig 3.**
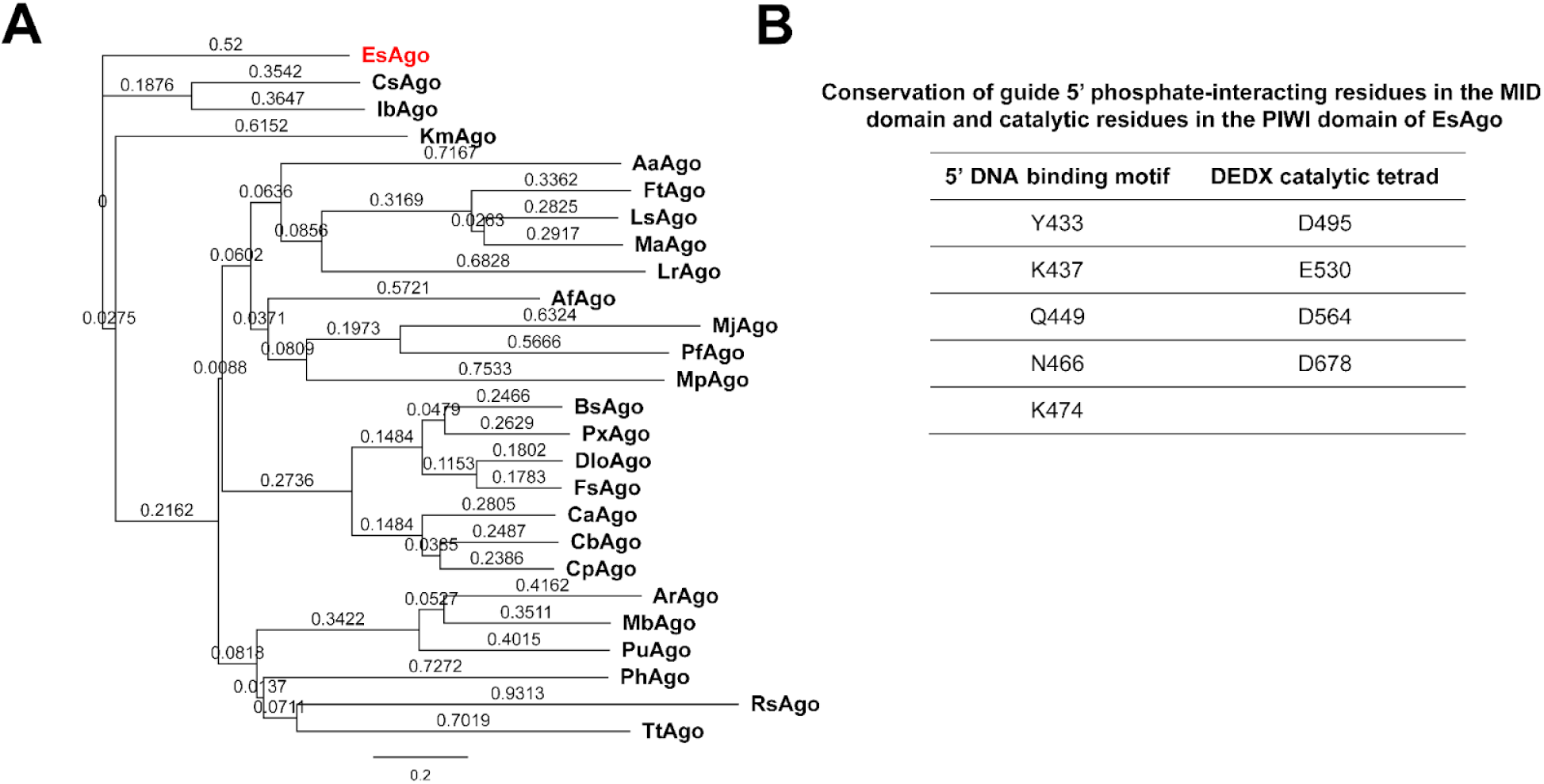
Sequence-structural analysis of EsAgo. (**A**) Neighbor-joining phylogenetic tree (global alignment, BLOSUM62 cost matrix, EsAgo as outgroup) of EsAgo with experimentally characterized pAgos from various genera. Branch labels display substitutions per site. Protein accession numbers are listed in **Supplementary Table 3**. (**B**) Conserved residues composing the YK-type and DEDX motifs of EsAgo.

A detailed view of the alignment between EsAgo and some of these characterized pAgos is presented in **Supplementary Fig. 2.** This revealed that EsAgo possesses a YKQXNK motif (i.e. YK-type DNA binding motif, where X can be any amino acid) in the MID domain similar to long pAgos that preferentially acquire 5’-P ssDNA guides, as well as a DEDD tetrad in the PIWI domain which suggests that EsAgo is a catalytically active nuclease (**Supplementary Fig. 2**). The specific residues which constitute these conserved motifs are summarized in **Fig. 3B**. Moreover, the presence of a full-length PAZ domain (**Supplementary Fig. 2**) [2], coupled with the presence of these two conserved motifs, suggest that EsAgo can be classified under the long-A clade of pAgos. Similarly, CsAgo, IbAgo, and KmAgo, are also classified as long-A pAgos and possess full-length PAZ domains, YK-type DNA binding motifs, and DEDX catalytic tetrads [2]. As these pAgos are described as DNA-guided DNA-interfering nucleases active at 37°C [38, 39, 40], we had thus speculated that EsAgo likewise exhibits nuclease activity and a preference for acquiring 5’-P ssDNA guides.

To determine possible structural characteristics of EsAgo, we constructed protein models using AlphaFold and PHYRE2 for *de novo* and homology-based prediction, respectively (**Supplementary Fig. 3A**). For the AlphaFold model, the vast majority of EsAgo was modeled at an acceptable confidence level (pLDDT > 70), though several distracting small stereochemical violations may have been introduced (**Supplementary Fig. 3B**) [36]. For PHYRE2, six solved Ago structures (eukaryotic and prokaryotic) with >90% sequence coverage and 100% confidence were collectively used as templates to generate a protein model: 4F1N [41], 6KR6 [42], 3HO1 [43], 3DLB [44], 5GUH [45], and 4OLB [46]. Based on both AlphaFold and PHYRE2 models, EsAgo is predicted to adopt a bilobal structure wherein the N and PAZ constitute one lobe and the MID and PIWI constitute the other, akin to all known full-length Agos (**Supplementary Fig. 3A**). Indeed, this bilobal arrangement is considered a defining characteristic of the tertiary structure adopted by long pAgos [2]. Additionally, the MID and PIWI of the AlphaFold model appear to adopt conformations similar to that of a Rossman-like fold and an RNase H fold, respectively, which is typical for these domains (**Supplementary Fig. 3C**) [1, 47, 48]. These were likewise observed in the PHYRE2 model (data not shown).

Taken together, these suggest that EsAgo is indeed a bona fide pAgo due to the presence of highly conserved residues in the functional domains and the adoption of an overall bilobal structure.

### EsAgo may shear its own expression vector in *E. coli*

As there have been a few documented cases of pAgos degrading their expression vectors in a heterologous host [15, 16, 49], we sought to functionally characterize EsAgo by probing this capacity for *in vivo* plasmid interference in *E. coli* as no genetic manipulation techniques are available for *E.* AB2. To qualitatively assess plasmid integrity in response to EsAgo WT expression, *EsAgo* was cloned into a T5-lacO-based expression vector, pQE-80L (subsequently generating pSML0001), and expressed in *E. coli* by way of IPTG induction, whereupon plasmids were extracted hourly then resolved on an agarose gel (**Fig. 4A**).

**Fig. 4.**
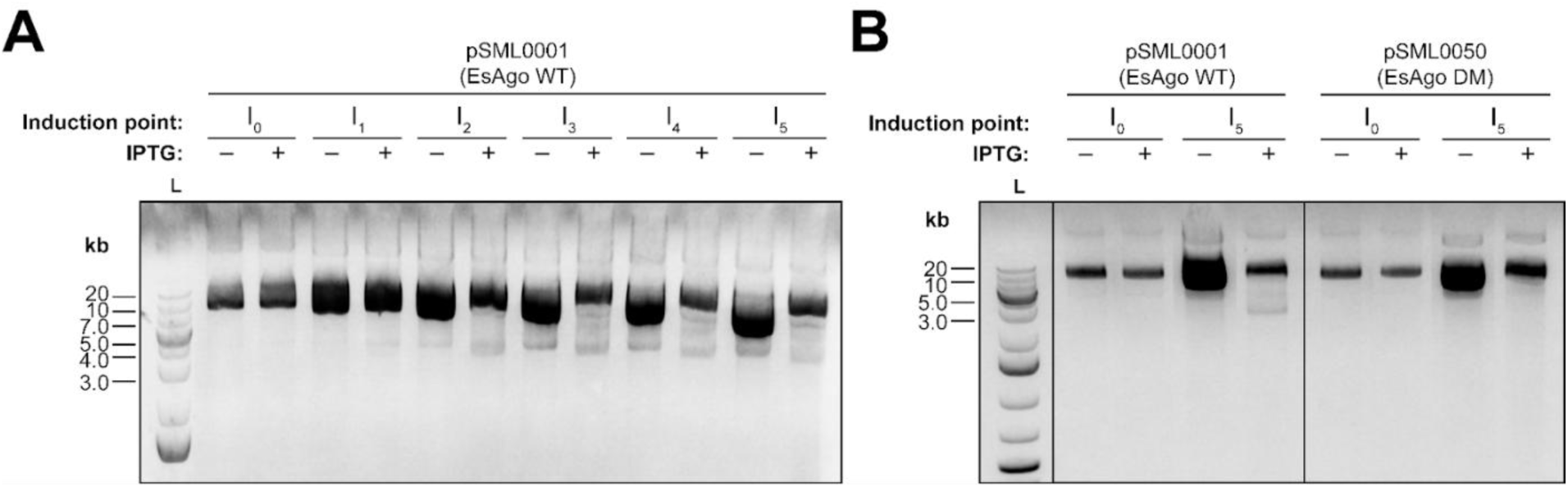
Plasmid minipreps of EsAgo expression vectors following IPTG induction in *E. coli*. **(A)** Plasmid quality decreases in response to longer periods of WT EsAgo overexpression. Cells were grown for a total of 8 hours with protein expression induced for a total of 5 hours once OD_600_ reached ∼0.6-0.8. Plasmids were extracted hourly following IPTG induction (I_0_-I_5_) and resolved through an agarose gel. Plasmids extracted from uninduced flasks without IPTG served as negative controls. (**B**) Plasmid shearing appears attenuated when the catalytic double mutant of EsAgo was overexpressed. Cells were grown and induced akin to those in (A). Plasmids were extracted from aliquots taken before (I_0_) and after 5 hours (I_5_) of induction then resolved on agarose gels. Plasmids from uninduced flasks without IPTG served as negative controls.

Here, plasmids extracted prior to IPTG induction appear to be of similar quality as the main ∼10 kb supercoiled bands are of similar concentration and the overall banding patterns appear consistent to one another (**Fig. 4A**, **I_0_**). However, plasmids from cells supplemented with IPTG appear to exhibit a “laddering” effect after 3 to 5 hours of induction as evidenced by the presence of additional bands, particularly at the ∼4 and ∼7 kb mark, which are notably absent among the uninduced cells (**Fig. 4A, I_3_-I_5_, lanes 7-12**). Moreover, the reduced concentration of the main ∼10 kb supercoiled band coupled with the appearance of a prominent ∼20 kb band may suggest an increase in linearized and/or open-circular plasmid conformations (**Fig. 4A, I_2_-I_5_, lanes 6, 8, 10, and 12**). These observations suggest that pSML0001 may be degraded and subsequently fragmented over time during EsAgo WT overexpression.

To determine if this plasmid degradation could be attributed to the nuclease activity of EsAgo WT, we likewise examined the quality of the vector expressing EsAgo DM (pSML0050) before and after a 5-hour induction period (**Fig. 4B**). Prior to the addition of IPTG, plasmid extracts appeared to be of similar yield and banding pattern between EsAgo WT and DM (**Fig. 4B, I_0_, lanes 1-2 and 5-6**). After 5 hours of induction, a reduction in concentration and a “laddering” effect of the expression vector of WT EsAgo was again observed (**Fig. 4B, lane 4**). Notably, plasmid “laddering” appeared to be attenuated following the expression of EsAgo DM—though plasmid concentration was reduced compared to the uninduced control (**Fig. 4B, lane 8**). We attempted to quantitatively examine this potential for plasmid interference by performing a cell viability assay wherein cells expressing EsAgo WT and DM were plated on media with increasing plasmid selection pressure (**Supplementary Fig. 4**). Assuming that EsAgo selectively degrades its expression vector under these conditions, we had thus expected to observe a significant reduction in bacterial viability in response to increasing plasmid selection. However, we had observed a general toxicity for both EsAgo WT and DM expression leading to cell death regardless of the media supplied (**Supplementary Fig. 4**). It is possible that cell death due to toxic expression may have contributed to reduced plasmid concentrations for EsAgo WT and DM following induction (**Fig. 4B, I_5_, lanes 4 and 8**). Nevertheless, the difference in banding patterns between EsAgo WT and DM expression vectors post-induction (**Fig. 4B, I_5_, compare lanes 4 and 8**) suggests that EsAgo may function as an active nuclease capable of targeting and degrading its own expression plasmid when expressed in *E. coli*.

### EsAgo requires a divalent cation for activity and degrades plasmids in a nonspecific guide-independent manner *in vitro*

Given that overexpression significantly reduced cell viability (**Supplementary Fig. 4**), this prevented us from further using *E. coli* as a cell model from which to probe the capacity of EsAgo for MGE-interference *in vivo*. This therefore necessitated studying EsAgo in a cell-free system *in vitro*, for which purified proteins were required. As we had determined that EsAgo was expressed in soluble (i.e. functional) form in *E. coli*, we purified N-terminally His_6_-tagged EsAgo WT and DM by FPLC (**Supplementary Fig. 5**). Attempts at isolating co-purifying short nucleic acid guides by way of PCI and TRIzol^TM^ extraction yielded empty lanes when resolved on PAGE gels (**Supplementary Fig. 6**), suggesting that EsAgo may purify free of associated guides akin to some pAgos [14, 40]. Despite this, guides are not a strict requirement for catalytic activity as some pAgos can degrade plasmid substrates in a random guide-independent manner [3, 16, 23, 24].

To determine if His_6_-EsAgo WT was functionally active *in vitro*, we incubated EsAgo with pQE-80L at 37°C in reaction buffer (see Methods for components) formulated with 1 mM of various divalent cations (**Fig. 5A**). Incubating EsAgo with Cu^2+^, Fe^2+^, Ni^2+^, Mg^2+^, and Mn^2+^ resulted in a noticeable reduction in plasmid concentration compared with controls (**Fig. 5A, compare lanes 1-4 with lanes 10, 12, 14, 16, and 18**). Plasmid degradation appeared most pronounced when EsAgo was supplied with Ni^2+^, Fe^2+^ and Mn^2+^ which resulted in about 79%, 94%, and 97% of supercoiled plasmids cleaved, respectively (**Supplementary Fig. 7**). Interestingly, supplying EsAgo with Cu^2+^ and Fe^2+^ resulted in increased intensities of the open-circular (OC) band concurrent with the reduction of the supercoiled band, thereby suggesting that plasmid conformations are relaxed (**Fig. 5A, lanes 10 and 12**). Minimal plasmid degradation was observed when EsAgo was incubated without any divalent cation (**Fig. 5A, lane 5**). Significantly, plasmid degradation was substantially attenuated when divalent cations were chelated with 10 mM EDTA as overall banding patterns and percent supercoiled plasmids cleaved appeared similar to controls (**Fig. 5A and Supplementary Fig. 7**). Collectively, these imply that EsAgo is capable of catalyzing DNA cleavage of plasmid substrates *in vitro* when incubated at 37°C and supplied with an appropriate divalent cation. To enable the efficient cleavage of DNA targets, the reaction buffer was formulated with a working concentration of 1 mM Mn^2+^ for all subsequent *in vitro* assays (**Supplementary Fig. 8**).

**Fig. 5.**
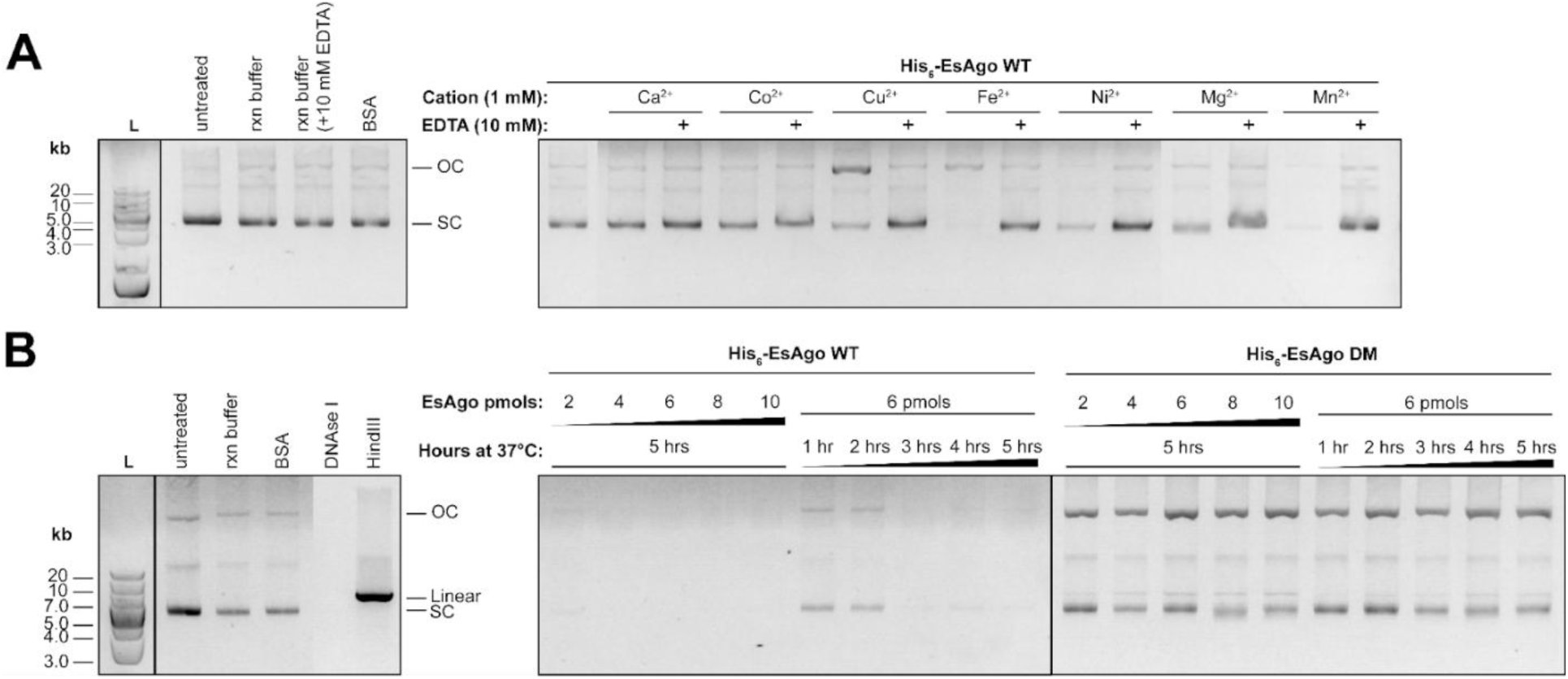
EsAgo randomly degrades pQE-80L when supplied with Mn^2+^ *in vitro*. (**A**) Analysis of the dependence of His_6_-EsAgo-mediated DNA cleavage on a divalent cation cofactor. Various divalent cations were supplied at 1 mM concentrations to 30 pmols of EsAgo and incubated with 500 ng of pQE-80L at 37°C for 5 hours. Plasmids incubated with 30 pmols of BSA served as a nonreactive protein negative control. (**B**) Substituting the D495 and D564 catalytic residues with alanines reduces the degree of nuclease activity exhibited by EsAgo. Plasmid quality in response to increasing amounts of protein or incubation time with EsAgo WT or DM was examined. Reactions consisted of 100 ng of plasmid incubated at 37°C with increasing amounts of protein for 5 hours or with 6 pmols of protein for 1-5 hours. Plasmids incubated with 10 pmols of BSA or 10 pmols of DNAse I served as negative and positive controls, respectively, for nuclease-mediated plasmid degradation. Plasmids digested with HindIII indicate linearized pQE-80L.

To further confirm if this *in vitro* plasmid degradation was indeed due to the catalytic activity of EsAgo, we investigated if EsAgo DM exhibits markedly decreased levels of this nuclease activity. To this end, pQE-80L was incubated for 5 hours at 37°C with increasing amounts of EsAgo WT or EsAgo DM, or for 1-5 hours with a fixed amount of protein (**Fig. 5B**). Incubating plasmids with EsAgo WT resulted in significant reductions in plasmid concentration across all setups (**Fig. 5B, lanes 6-15**) with substantial amounts of supercoiled plasmids being cleaved (**Supplementary Fig. 7**). Incubation with ≥6 pmols of EsAgo WT for 5 hours or 6 pmols of EsAgo WT for ≥3 hours resulted in 100 ng of pQE-80L being almost fully digested as only faint bands, if any, remained on the AGE gel (**Fig. 5B, lanes 8-10 and 13-15**) with ∼89.5-99.5% of supercoiled plasmids cleaved (**Supplementary Fig. 7**). Additionally, the lack of recurring banding patterns from EsAgo WT digestion (**Fig. 5B, lanes 6-15**), the destruction of all plasmid conformations to a degree comparable with DNAse I digestion (**Fig. 5B, lane 4**), and the apparent lack of associated co-purifying guides (**Supplementary Fig. 6**) collectively suggests that EsAgo degrades plasmids in a random guide-independent manner akin to other apo-pAgos [3].

Strikingly, this high degree of degradation was decreased across all EsAgo DM reactions with no instances of near-complete plasmid digestion occurring (**Fig. 5B, lanes 16-25**). In contrast to EsAgo WT, EsAgo DM-mediated plasmid degradation does not appear to increase in severity with greater protein dosages or longer incubation times as evidenced by the generally consistent band intensities between reactions (**Fig. 5B, lanes 16-25**). It should be noted that while an increase in supercoiled plasmids cleaved in response to greater EsAgo DM dosages (∼25-60% cleavage) and longer incubation times (∼38-66% cleavage) was observed, this was coupled with an evident increase in open-circular and linearized plasmids (**Fig. 5B, lanes 16-25**) (**Supplementary Fig. 7**). This suggests that EsAgo DM retains some residual nuclease activity and may be nicking and cleaving supercoiled pQE-80L into open-circular and linearized plasmid conformations, respectively.

### Supplying EsAgo with forward and reverse guides results in plasmid linearization

As we had determined that EsAgo is catalytically active *in vitro*, our final objective was to probe the capacity of EsAgo for targeting specific nucleic acid sequences using complementary guides—a fundamental feature shared by all Ago proteins [2]. We designed synthetic ssDNA oligos 21 nt in length with 5’-phosphate groups targeting a region with low GC content (32% GC) within the *lacI* gene of pQE-80L (**Fig. 6A**) as pAgos have been noted to cleave GC-rich DNA inefficiently (**see Supplementary Table 4 for all guides used in this study**) [17, 23, 40]. Forward and reverse guides (one EsAgo-guide complex per strand) were arrayed to generate staggered 5 nt-long 3’-ssDNA overhangs based on the typical pAgo cleavage site between the 10th and 11th nucleotide from the 5’ end of the guide (**Fig. 6A, black triangles**) [17]. This was done as the spacing between cleavage sites may significantly affect the efficiency of generating double-stranded breaks [17] with Vaiskunaite *et al*. [38] reporting that this type of cut is most efficient for guided dsDNA cleavage by CbAgo. Additionally, CbAgo-mediated guided dsDNA cleavage has been reported to depend in large part on an appropriate molar ratio between guide and pAgo [38]. When we supplied EsAgo WT with various concentrations of *lacI*-targeting guides, we determined that a guide:EsAgo ratio of 1:1 and 1:3 was most efficient for the forward and reverse guides, respectively (**Supplementary Fig. 9**). Interestingly, minimal plasmid degradation was observed when EsAgo was supplied with various doses of guides (**Supplementary Fig. 9**) in contrast to the nonspecific guide-independent nuclease activity seen in **Fig. 5B** (**lanes 6-15**). Indeed, following the incubation of pQE-80L with EsAgo WT-guide complexes for extended periods of 20 and 40 hours, plasmids remained largely intact while those incubated with guide-free EsAgo WT were completely destroyed (**Supplementary Fig. 10**). These indicate that the random DNA “chopping” activity exhibited by apo-EsAgo is inhibited by the presence of guides.

**Fig. 6.**
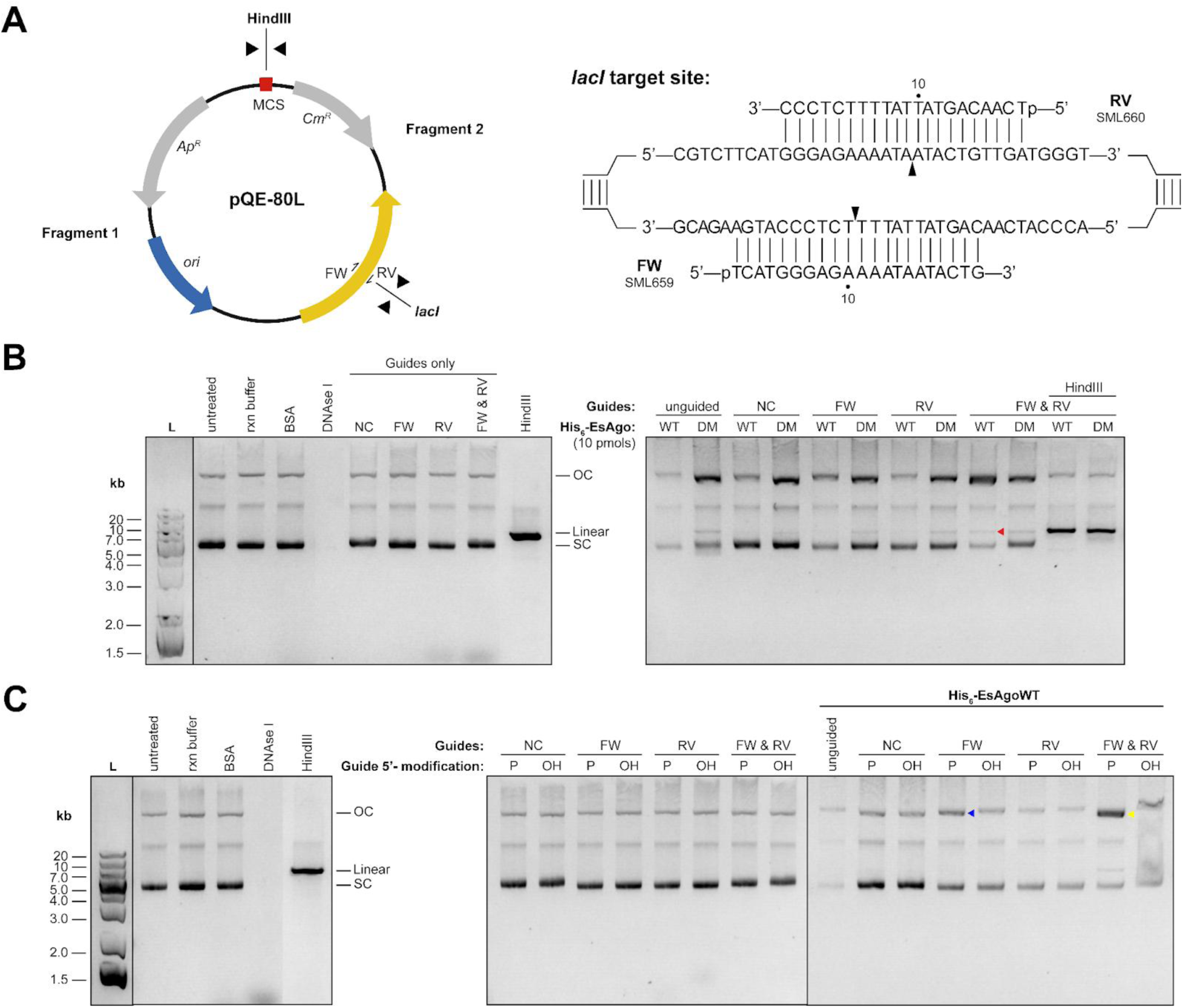
Analysis of the programmable cleavage of plasmid dsDNA by EsAgo. (**A**) Synthetic 21 nt-long 5’-P ssDNA guides were used to target the *lacI* gene (32% GC) of pQE-80L. Forward (FW) and reverse (RV) guides were arrayed to generate 5 nt-long staggered 3’-ssDNA overhangs. Black triangles indicate the expected cleavage site of each guide. Subsequent digestion with HindIII was expected to produce two DNA fragments of known length. (**B**) Guides were loaded in two half-reactions wherein 5 pmols of His_6_-EsAgo WT or His_6_-EsAgo DM were incubated at 37°C for 30 minutes with forward or reverse guides. Guides were loaded onto EsAgo at a 1:1 molar ratio while the reverse guide was loaded at a ratio of 1 guide:3 EsAgo. Half-reactions were mixed and incubated at 37°C for 5 hours with 100 ng of pQE-80L. The red arrow indicates an increase in linearized pQE-80L when incubated with both EsAgo WT-FW and EsAgo WT-RV complexes. Reactions were subsequently digested with HindIII for 1 hour. (**C**) Comparison of EsAgo WT-mediated plasmid cleavage with 5’-P or 5’-OH *lacI* guides. Reactions were set up akin to those in (A). The blue arrow indicates an increase in open-circular plasmids when EsAgo WT was supplied with 5’-P *lacI* FW while the yellow arrow indicates a drastic increase in open-circular band intensity with 5’-P *lacI* FW and RV.

To facilitate the programmable cleavage of dsDNA, FW and RV guides were loaded onto WT and DM EsAgo and incubated with pQE-80L for 5 hours at 37°C (**Fig**. **6B**). To confirm if pQE-80L was cleaved at the intended site by EsAgo-guide complexes, plasmids were subsequently digested with HindIII for 1 hour as double-stranded breaks sustained within the MCS and at the targeted *lacI* sequence would yield two DNA fragments of known size (**Fig. 6A**). Oddly, the residual catalytic activity seen in guide-free EsAgo DM (**Fig. 5B, lanes 16-25**) appeared unaffected by the presence of guides (**Fig. 6B, compare lane 11 with lanes 13, 15, 17, and 19**) as increases in open-circular and linearized plasmids were observed regardless of the presence, type, or combination of guides supplied (also observed in **Supplementary Fig. 10**) (**Supplementary Fig. 11A and B**).

On the other hand, plasmids incubated with EsAgo WT-guide complexes exhibited changes in conformation depending on the guide provided (**Fig. 6B**). When EsAgo WT was supplied with a “scrambled” non-complementary (NC) control guide, pQE-80L band profiles on AGE gels were highly similar to plasmid controls incubated in reaction buffer only (**Figure 6B, compare lanes 2 and 12**) with only a minimal ∼0.27-fold increase in open-circular plasmids measured (**Supplementary Fig. 11B**). When supplied with individual forward and reverse guides, EsAgo WT-FW complexes yielded substantial ∼1.12-fold increases in open-circular plasmids, whereas no measurable increases in the open-circular state were observed among EsAgo WT-RV plasmid digests relative to controls (**Supplementary Fig. 11B**). This suggests that single-stranded breaks sustained at the site targeted by the FW guide may be nicking supercoiled pQE-80L into an open-circular form. Expectedly, no plasmids were linearized following incubation with either EsAgo WT-FW or EsAgo WT-RV complexes alone (**Fig. 6B**) (**Supplementary Fig. 11A**). However, when EsAgo WT was supplied with both forward and reverse guides, this consistently resulted in the appearance of a faint ∼8-9 kb band (**Fig. 6B, lane 18, red arrow**) corresponding to an average of ∼5.67% of plasmids being linearized relative to controls (**Supplementary Fig. 11A**). Notably, this modest increase in linearized plasmid was coupled with a drastic ∼3.73-fold increase in open-circular plasmids which suggests that plasmids are cleaved and nicked by EsAgo in a guide-dependent manner (**Supplementary Fig. 11B**). Unfortunately, the subsequent digestion of these plasmids with HindIII did not yield the expected ∼1.6 kb and ∼3.1 kb fragments, thereby making it unclear if EsAgo-mediated dsDNA cleavage occurred at the intended sequence (**Fig. 6B, lane 20**).

As an alternative means of probing EsAgo-mediated programmable dsDNA cleavage, we supplied EsAgo WT with guides bearing 5’-OH groups (but were otherwise identical in sequence) as DNA cleavage efficiency has been noted to decrease when using non-phosphorylated guides [2]. Reactions were set up identically to those in **Fig. 6B** and the corresponding changes in plasmid conformations observed (**Fig. 6C**). Here, the random nuclease activity exhibited by unguided EsAgo WT was attenuated when supplied with either 5’-P or 5’-OH guides, particularly when NC guides were provided (**Fig. 6C**, **lanes 15-16**). Notably, a ∼1.92-fold increase in open-circular plasmids was observed in response to loading EsAgo WT with 5’-P forward guides, whereas loading 5’-OH forward guides resulted in a moderate ∼0.29-fold increase in open-circular plasmids (**Fig. 6C, lane 17, blue arrow**) (**Supplementary Fig. 11C**). Oddly, EsAgo WT supplied with 5’-P or 5’-OH reverse guides yielded a ∼0.66-fold and a ∼0.62-fold reduction in open-circular plasmids, respectively (**Supplementary Fig. 11C**). Significantly, plasmids exhibited a ∼1.92-fold increase in open circular plasmids when EsAgo WT was loaded with both 5’-P forward and reverse guides which was notably absent when supplied with both 5’-OH guides (**Fig. 6C, lane 21, yellow arrow**) (**Supplementary Fig. 11C**). Although modest amounts of plasmids were linearized when both 5’-P *lacI* guides were supplied (consisted with our observations in **Fig. 6B**), DNA smearing makes it unclear if this also occurred when EsAgo WT was provided with 5’-OH forward and reverse guides (**Fig. 6C, lane 22**).

Thus, the changes in the conformation of pQE-80L we had repeatedly observed when EsAgo was supplied with 5’-P forward and reverse guides suggests that EsAgo may be capable of catalyzing the cleavage of dsDNA in a programmable sequence-specific manner. As such, this carries with it implications in the role of EsAgo as a defense system and its potential utilization as a biotechnological tool.

## DISCUSSION

Bacterial evolution is in large part driven by the constant horizontal uptake of foreign genetic elements. While these can confer beneficial traits (e.g. antibiotic and toxin resistance) onto their bacterial hosts, these are fundamentally selfish genetic entities [8] and are often parasitic in nature [27]. As such, bacteria have evolved an “immune system” comprising a vast arsenal of sophisticated defense systems typically organized into genomic “defense islands” [28, 30]. Accordingly, genes with unknown function residing within these “defense islands” are hypothesized to provide protection against MGEs [30]. In the case of *E.* AB2, our search for novel defense systems had led us to discover a “defense island” containing genes encoding putative type II TA, type II Wadjet, type I RM, and DISARM systems clustered upstream of *EsAgo* and other closely related *pAgos* from various *Exiguobacterium* species (**Fig. 2**). It follows that the proximity of *EsAgo* (and other *Exiguobacterium* pAgos) to known defense systems—a common phenomenon observed among pAgos [50], strongly suggests a role in host defense [10].

The most commonly encoded defenses are those that directly antagonize intracellular foreign nucleic acids in a sequence-specific manner [27]. These include RM systems, CRISPR-Cas systems, and long-A pAgos [27]. Our computational analysis revealed that *EsAgo* indeed encodes a bona fide long-A pAgo as EsAgo possesses the requisite motifs and is predicted to exhibit the structural characteristics typical of this pAgo clade (**Fig. 3**). Agreeably, the extensive bioinformatics analysis on pAgos conducted by Ryazanksy *et al*. [2] lists EsAgo as a member of the long-A clade.

As a long-A pAgo, EsAgo is a catalytically active enzyme capable of degrading DNA substrates. However, we could not examine where EsAgo-mediated nucleic acid interference is directed within *E.* AB2 due to the absence of genetic manipulation techniques for *Exiguobacterium* species. We instead investigated if EsAgo degrades plasmids *in vivo* akin to some pAgos heterologously expressed in *E. coli* (**Fig. 4**) [15, 49]. Although plasmid “laddering” in response to EsAgo WT expression was attenuated during the expression of EsAgo DM (**Fig. 4B**), the toxic effects of expressing these proteins in *E. coli* (**Supplementary Fig. 4**) makes it unclear if the observed plasmid degradation was solely due to EsAgo-mediated plasmid interference, significantly reduced cell viability, potential off-target effects (e.g. collateral genomic degradation by EsAgo), or any combination of these factors. Indeed, it has been reported that pAgos are poorly expressed in *E. coli*—the cause of which has been ascribed to a general toxicity of pAgos during expression [49]. To improve expression, most studies typically induce cultures at lower temperatures of 16-18°C [13, 14, 17, 40, 49]. It is possible that our expression of EsAgo at 37°C may have exacerbated this toxicity. Notably, the expression of a catalytic double mutant of the closely related EmaAgo (89.46% identity) at 30°C was also reported as toxic to *E. coli* [18]. Significantly, this toxicity prevented us from conducting phage infection and plasmid transformation efficiency assays on cells expressing EsAgo. Thus, it remains unclear if EsAgo can selectively interfere with MGEs in *E. coli*.

Similar to most studies, we had instead characterized EsAgo-mediated plasmid degradation *in vitro*. Here we determined that EsAgo utilizes Mn^2+^ as a cofactor and randomly “chops” plasmids in the absence of available guide strands (**Fig. 5**). In contrast to most characterized long-A pAgos which perform guide-independent plasmid “chopping” slowly [3, 14, 16, 23, 24, 40, 52], apo-EsAgo appears to process plasmids efficiently with substantial, and at times complete, degradation of pQE-80L repeatedly observed (**Fig. 5B and Supplementary Fig. 7B**). While the majority of these *in vitro* assays were performed on supercoiled pQE-80L, we note that unguided EsAgo was capable of randomly degrading other DNA substrates such as the circular ssDNA genome of M13 phage and pre-linearized pQE-80L (**Supplementary Fig. 12**), though testing on RNA targets remains open to future investigation.

Supercoiled plasmid degradation by guide-free EsAgo was typically extensive while EsAgo-guide complexes appeared to nick or cleave supercoiled pQE-80L into open-circular or linearized states, respectively. This suggests that EsAgo may preferentially target supercoiled plasmids similar to TtAgo [13], CbAgo [17], and KmAgo [40]. It is hypothesized that the supercoiled topology facilitates local DNA melting (particularly at AT-rich sequences), thereby providing regions accessible to pAgo targeting and cleavage [13]. Speculatively, pAgos may prioritize targeting foreign supercoiled plasmids as transcription levels from open-circular and linearized plasmids are decreased due to having sustained DNA breaks [13, 52]. Indeed, the conversion of supercoiled pQE-80L into open-circular and linearized forms by EsAgo DM, despite limited catalytic activity, appears to support this hypothesis (**Fig. 5B and Supplementary Fig. 7B**). Simultaneously, this implies that even a catalytically impaired EsAgo may provide host protection. Intriguingly, DEDX catalytic double mutants of *Natronobacterium gregoryi* Ago (NgAgo) [53], CbAgo [19], and EmaAgo (D494A, D564A) [18] appear to retain some function as well [49]. Notably, CbAgo DM and EmaAgo DM were capable of protecting *E. coli* from phage infection (though less efficiently than their respective WT counterparts) presumably because pAgo binding interferes with phage replication and/or repair [18, 19]. Moreover, mutations at the DEDX motif may alter guide acquisition as CbAgo DM was noted to exhibit weaker guide binding (versus WT) which may explain why the limited catalytic activity of EsAgo DM was not attenuated by ssDNA guides (**Fig. 6B**) (**Supplementary Figs. 10, 11A and B**) [19, 49]. On a related note, the random nuclease activity of LrAgo [19] and EmaAgo [18] was largely unaffected by the addition of ssDNA guides which suggests that other factors may be needed to suppress the guide-independent “chopping” of pAgos. Collectively, these indicate that EsAgo may interfere with MGEs via direct nucleolytic degradation or possibly through non-catalytic binding.

The use of short nucleic acid guide strands represents a major mechanism by which pAgos distinguish “non-self” genetic elements prior to mounting an appropriate effector response (e.g. specific cleavage, non-catalytic binding, etc.) [3, 54]. Indeed, the attenuation of indiscriminate plasmid “chopping” when synthetic ssDNA guides were supplied (**Fig. 6**) (**Supplementary Figs. 9** and **10**) suggests that EsAgo switches to a specific mode of action so as to prevent collateral damage of “self” genetic elements [3, 55]. Unfortunately, we could not determine the target preferences of EsAgo as we did not detect any co-purifying short nucleic acid guides (**Supplementary Fig. 6**). Though not all pAgos co-purify with bound guides following expression in *E. coli*, EmaAgo co-purifies with ssDNA guides enriched in phage and plasmid sequences [18]. This suggests that discrepancies in pAgo expression and purification protocols, rather than the absence of necessary guide-loading factors from *E.* AB2 (if any), may have resulted in guide-free EsAgo extracts. Indeed, two independent studies by Liu *et al*. [40] and Kropocheva *et al*. [56] report acquiring empty and guide-loaded KmAgo extracts, respectively, presumably for these reasons. Thus, it remains unclear if EsAgo, in its capacity as a defense system, preferentially sources guides from plasmids akin to other long-A pAgos heterologously expressed in *E. coli* [17, 18]. It should be noted, however, that such analyses may not accurately represent where pAgos source guides under native expression levels in their natural contexts [3]. Thus, future efforts should also be directed at examining EsAgo guide biogenesis pathways and other associated protein networks within its natural host if possible.

While it is likely that EsAgo preferentially utilizes 5’-P ssDNA guides to target and cleave MGEs akin to other long-A pAgos, it remains ambiguous if plasmids were nicked or cleaved in a sequence-specific manner by EsAgo-guide complexes (**Fig. 6**). When supplied with one guide, EsAgo-FW complexes consistently yielded marked increases in nicked open-circular plasmids from supercoiled pQE-80L, whereas EsAgo-RV complexes inconsistently produced modest increases in open-circular plasmids. It is unclear why other guides appeared non-functional or otherwise exhibited reduced functionality (**Supplementary Fig. 9**). Interestingly, discrepancies in the cleavage efficiencies of forward and reverse guides are also seen in some pAgos such as *Clostridium perfringens* Ago [39] and KmAgo [40]. Possible explanations include potential nucleotide biases (which we could not account for without co-purifying guides) and pAgo sensitivity to certain molar ratios of guides, among other factors [15, 24, 38]. Additionally, the limited accessibility of the DNA duplex at 37°C may inhibit cleavage by EsAgo-guide complexes as observed in other mesophilic pAgos [13, 38]. Within *E.* AB2, EsAgo may rely on associated helicases, such as the putative *drmA* helicase (**Fig. 1**), to access specific sequences on foreign dsDNA at moderate temperatures [19, 38]. Nevertheless, the consistent, albeit inefficient, linearization of pQE-80L when 5’-P FW and RV guides were supplied to EsAgo suggests that dsDNA cleavage may have occurred at the targeted *lacI* sequence (**Fig. 6**). However, this could not be confirmed as the expected DNA restriction fragments following HindIII digestion were not detected via AGE (**Fig. 6**). Moreover, guided cleavage assays on pre-linearized pQE-80L and the ssDNA genome of M13 phage also did not yield the fragments of interest (**Supplementary Fig. 12**). Thus, site-specific cleavage of DNA by EsAgo, while certainly a possibility, remains an open question for future studies to confirm.

## CONCLUSION

As nucleic acid-guided nucleic acid-targeting nucleases, pAgos have long been hypothesized as components of bacterial immunity against phages and other MGEs. By way of a computational analysis, we determined that EsAgo constitutes a bona-fide full-length pAgo of the long-A clade on the basis of its immediate genomic neighborhood, predicted structural features, and possession of the requisite motifs for catalytic activity and DNA guide binding. When heterologously overexpressed in *E. coli*, this may have resulted in EsAgo-mediated plasmid interference against its expression vector. Biochemical testing *in vitro* revealed that EsAgo, when supplied with an appropriate divalent cation cofactor, is a catalytically active DNA nuclease capable of efficiently degrading plasmids in a random guide-independent manner. Moreover, providing EsAgo with synthetic 5’-P ssDNA guides attenuates this random nuclease activity and, simultaneously, targeting pQE-80L with EsAgo-guide pair complexes may have resulted in the sequence-specific cleavage of dsDNA at 37°C. Thus, it is likely that EsAgo interferes with foreign DNA in both a guide-independent and a guide-dependent manner within *E.* AB2, though more *in vivo* data is needed. Lastly, the programmable cleavage of dsDNA at moderate temperatures by EsAgo for practical use remains open to investigation for which this study provides foundational data to be built upon.

## ABBREVIATIONS

5’-P: 5’-Phosphorylated
5’-OH: 5’-Hydroxylated
Abi: Abortive infection
Ago: Argonaute
Ap: Ampicillin
BREX: Bacteriophage Exclusion
DISARM: Defense island system associated with restriction-modification
DM: Double mutant
eAgo: Eukaryotic Ago
FW: Forward
HGT: Horizontal gene transfer
IPTG: Isopropyl β-D-1-thiogalactopyranoside
LB: Luria-Bertani Broth
LBA: LB agar
NC: Non-complementary
OC: Open-circular
OD600: Optical density at 600 nm
Oligos: Oligonucleotides
PADLOC: Prokaryotic Antiviral Defense Locator
pAgo: Prokaryotic Ago
Phages: Bacteriophages
RM: Restriction-modification
RNAi: RNA interference
RV: Reverse
TA: Toxin-antitoxin
TSB: Trypic Soy
SC: Supercoiled
SD: Standard deviation
WT: Wild type
EsAgo: *Exiguobacterium* sp. AB2 Ago
CbAgo: *Clostridium butyricum* Ago
CsAgo: *Clostridium sartagoforme* Ago
EmaAgo: *Exiguobacterium marinum* Ago
IbAgo: *Intestinibacter bartlettii* Ago
KmAgo: *Kurthia massiliensis* Ago
LrAgo: *Limnothrix rosea* Ago
MjAgo: *Methanocaldococcus janaschii* Ago
NgAgo: *Natronobacterium gregoryi* Ago
PfAgo: *Pyrococcus furiosus* Ago
RsAgo: *Rhodobacter sphaeroides* Ago
TtAgo: *Thermus thermophilus* Ago

