## Supplementary Table 1 for "Random guide-independent DNA cleavage from the Argonaute of *Exiguobacterium* sp. AB2"

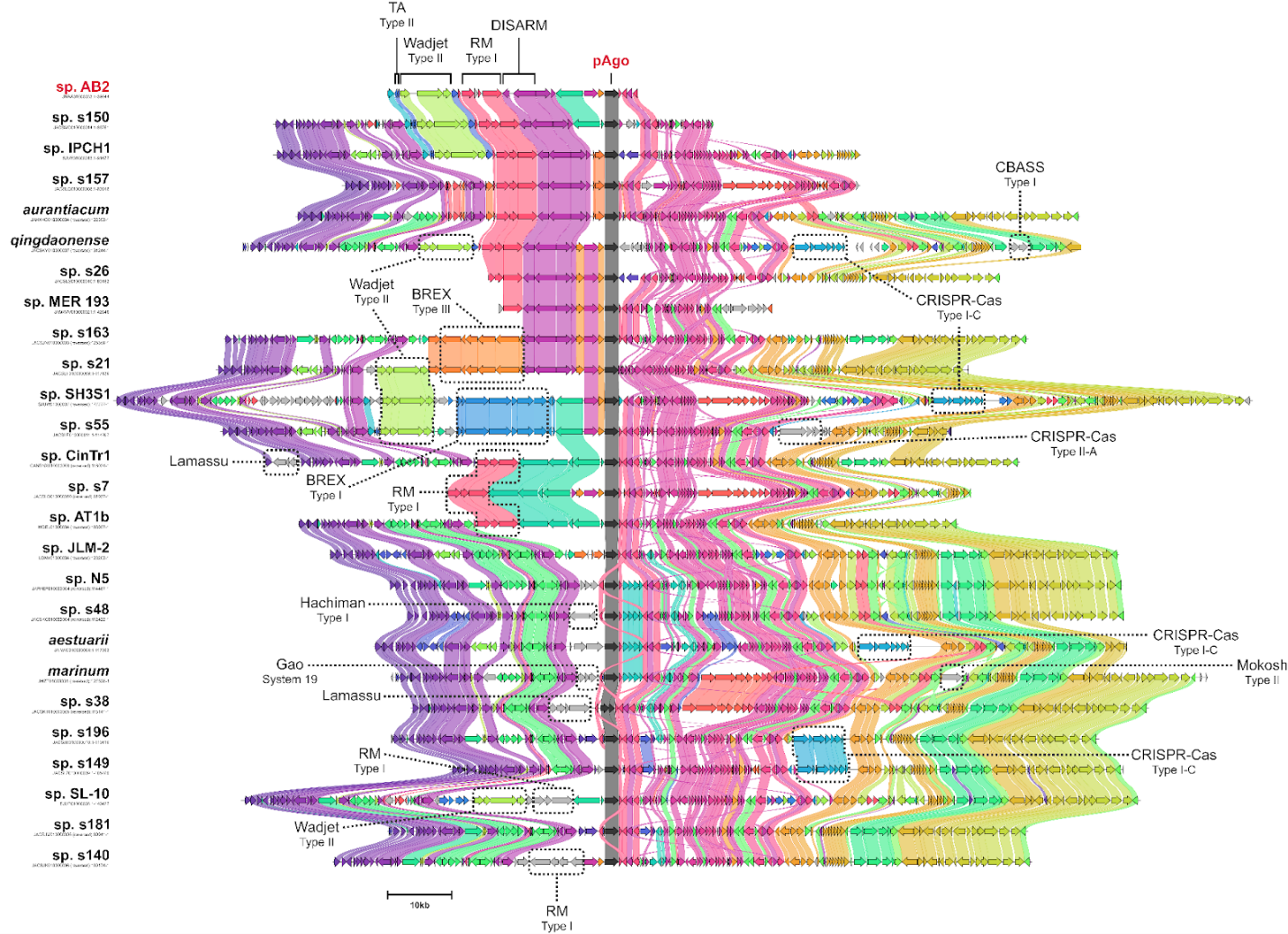

**Supplementary Fig. 1.** **Genomic neighborhoods of *Exiguobacterium* pAgos (extended).** Multiple genome alignment of contigs containing *Exiguobacterium* pAgos. The alignment is centered around the pAgo gene (dark gray arrows) while other defense systems identified by PADLOC are highlighted. NCBI accession numbers of the contigs used in the alignment are displayed under the *Exiguobacterium* species it belongs to and are listed in **Supplementary Table 1**. Contig alignment and gene cluster similarity were generated using clinker. Note: the contig derived from *E. marinum* is cropped due to its length. Additional defenses such as a type II RM, type IV, RM, and a VSPR were identified by PADLOC further downstream of the *E. marinum* contig (see **Supplementary Table 2**).

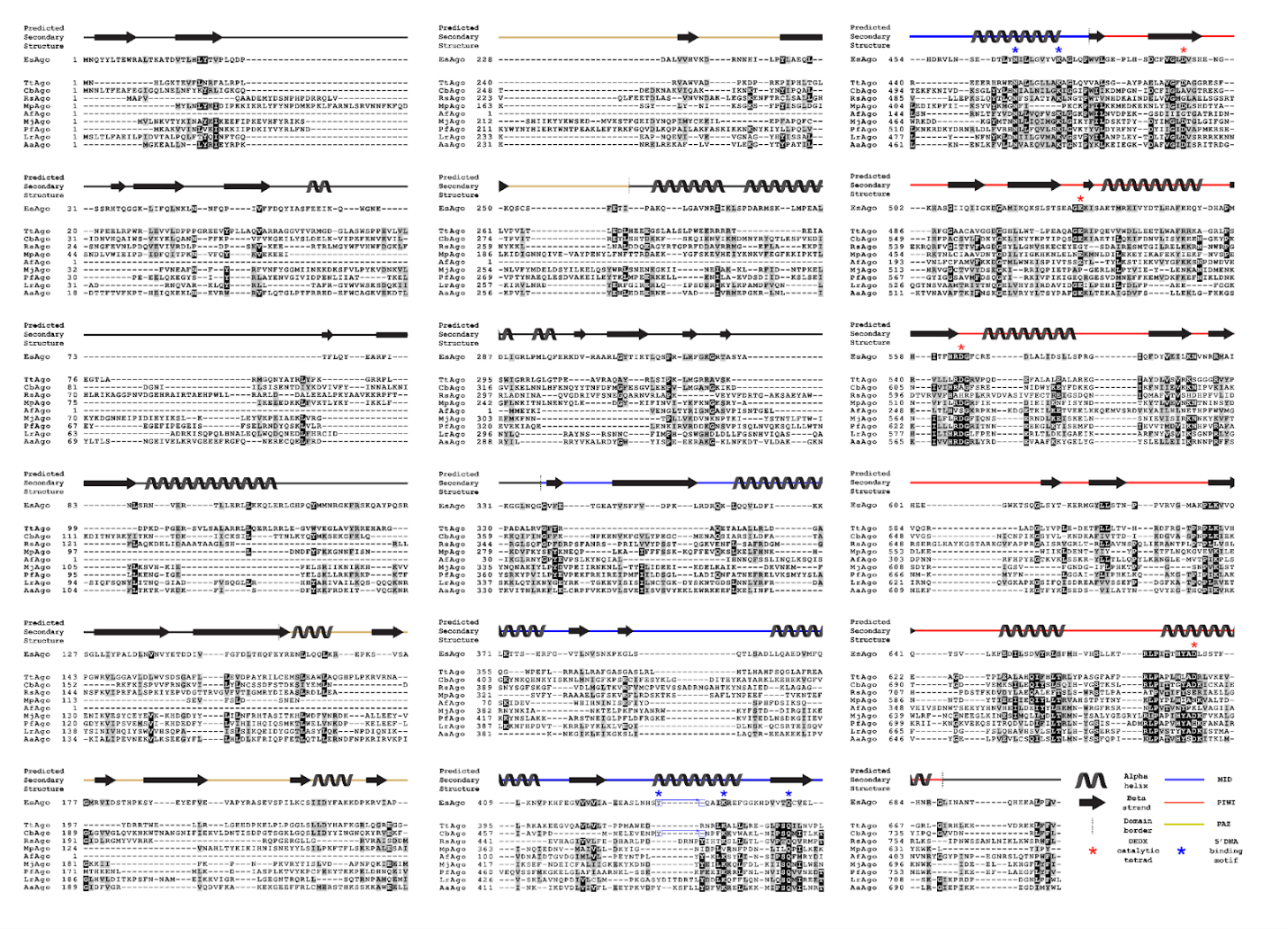

**Supplementary Fig. 2.** **Multiple sequence alignment of EsAgo with characterized pAgos from various bacterial genera.** Black and gray shading represent identical and similar residues, respectively. Alignment was generated using the ClustalO algorithm. PHYRE2 predicted secondary structures of EsAgo are displayed. Borders of the MID (blue), PAZ (yellow), and PIWI (red) domains of EsAgo were taken from the predictions of Ryazansky et al. [2]. Note: CbAgo sequence is a catalytic double mutant (D541A, D611A).

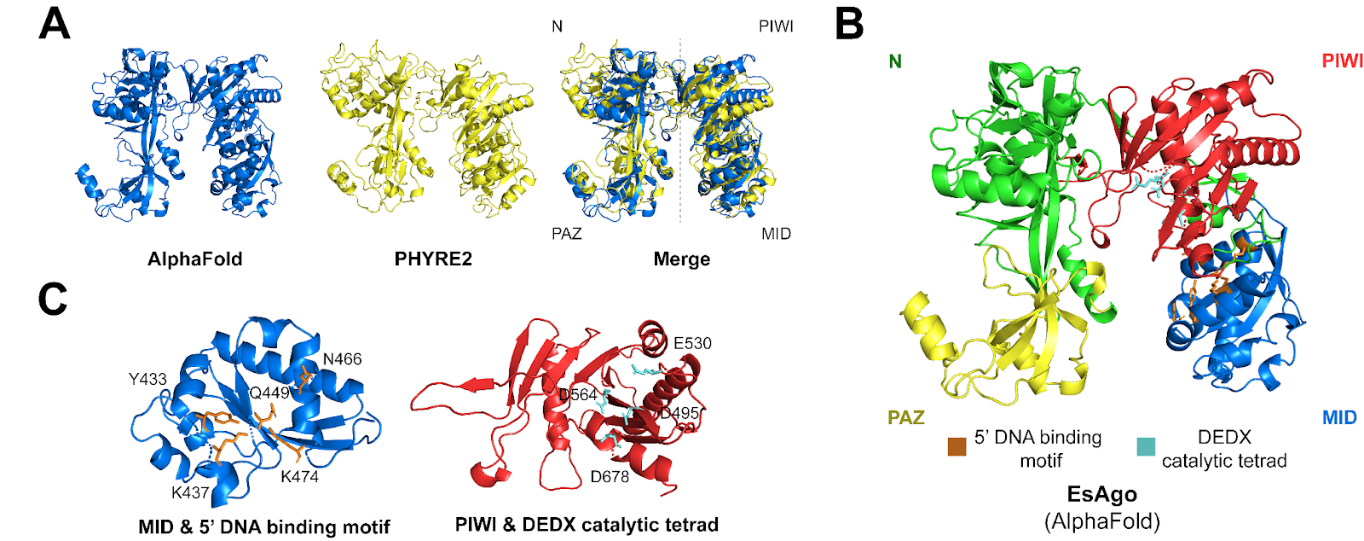

**Supplementary Fig. 3. Protein modeling of EsAgo.** (**A**) Protein models were generated using AlphaFold and PHYRE2. Arrows represent β-sheets while helices represent α-helices. (**B**) AlphaFold model of EsAgo with the PAZ, MID, and PIWI domains colored yellow, blue, and red, respectively. Green portions represent regions that are poorly conserved among pAgos such as the N-terminal domain. Domain borders were taken from Ryazansky *et al*. [2]. (**C**) Close-up views of the predicted AlphaFold MID and PIWI domains of EsAgo detailing the YK-type DNA binding motif and DEDX catalytic tetrad, respectively.

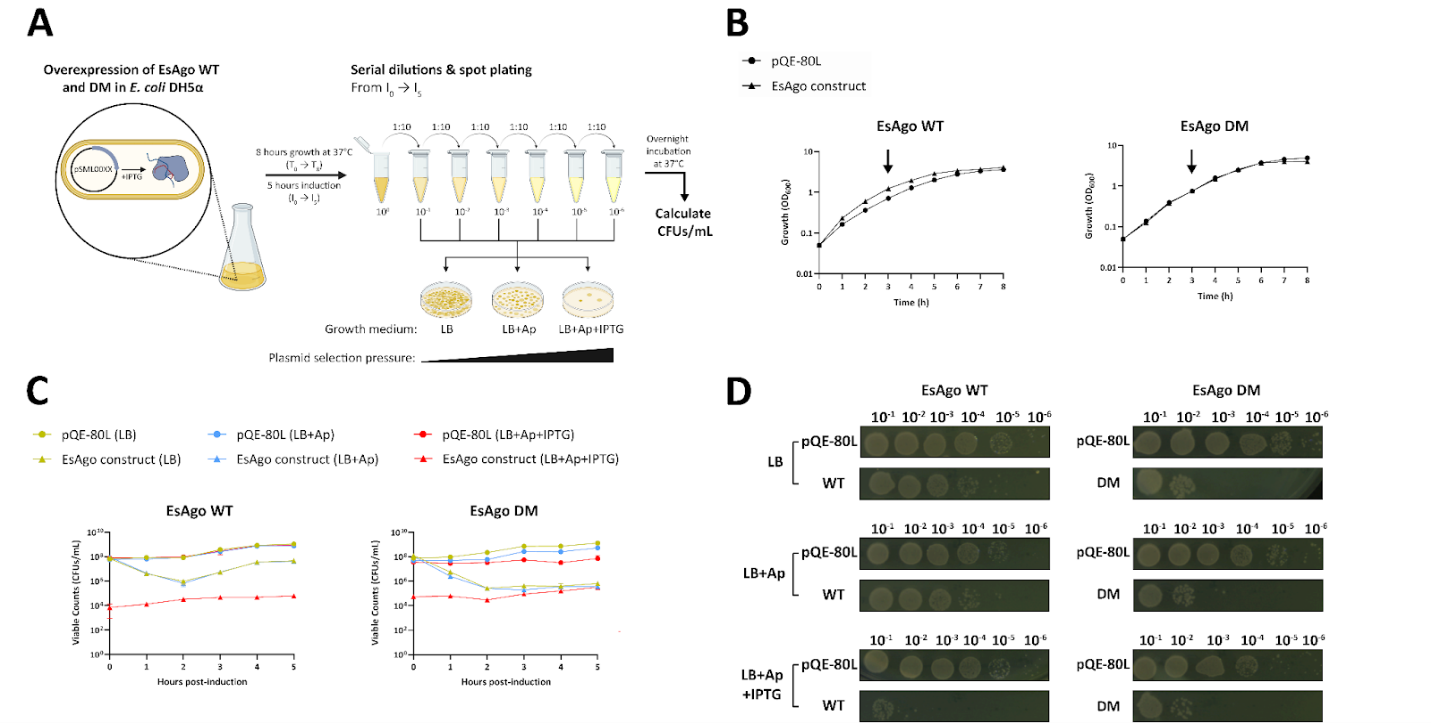

**Supplementary Fig. 4. Overexpressing EsAgo and its catalytic double mutant reduces cell viability in *E. coli*.** (**A**) Schematic diagram illustrating the experimental procedure of the cell viability assays performed. (**B**) Growth measurements of *E. coli* cells overexpressing WT or DM EsAgo constructs (triangles) or bearing empty pQE-80L plasmids (negative controls) (circles) over an 8-hour period. Overexpression of EsAgo constructs was induced with IPTG when OD_600_ reached ~0.6-0.8 (black arrow) and lasted for 5 hours. Aliquots were taken hourly from the point of induction (I_0_) until the end of the growth period (I_5_), spot plated, and used to determine bacterial viable counts. (**C**) Bacterial viable counts (measured as CFUs/mL) of cells overexpressing EsAgo constructs following 1-5 hours of induction in liquid media and overnight growth on LBA (yellow), LBA with Ap (blue), or LBA with Ap and IPTG (red). Error bars indicate SD of three technical replicates (n=3). (**D**) Representative images of the resulting *E. coli* colonies grown following a 5-hour induction period (I_5_) with IPTG, serial dilution and spot-plating onto various media, and overnight incubation at 37°C. Bacterial cells bearing empty pQE-80L plasmids grown under similar conditions were used as negative controls. Each spot is comprised of 10 µL of bacterial culture.

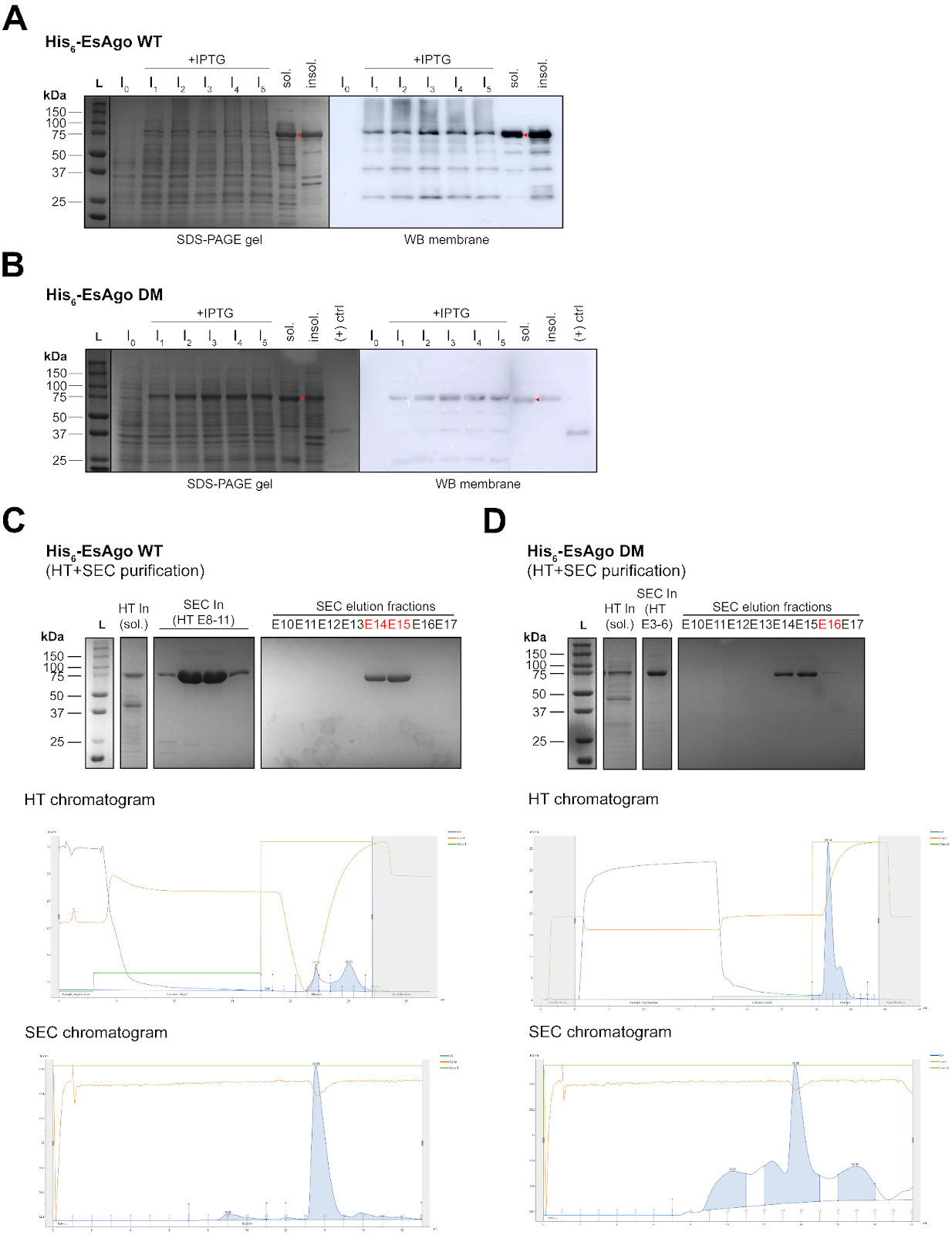

**Supplementary Fig. 5. Expression and purification of EsAgo from *E. coli*.** His_6_-tagged EsAgo (**A**) WT and (**B**) DM are expressed in both soluble and insoluble form in *E. coli* DH5ɑ. Cells were lysed hourly during His_6_-EsAgo expression (I_0_-I_5_) with total protein content resolved on 12% SDS-PAGE gels stained with Coomassie. Lysate taken at 5 hours post-induction (I_5_) was further separated by centrifugation into soluble (sol) and insoluble (insol) protein fractions. Western blots targeting His_6_-tags were performed in parallel to confirm protein expression. Red arrows indicate soluble full-length His_6_-EsAgo WT and DM. A ~40 kDa His_6_-tagged protein served as a positive control for western blot. Soluble (**C**) His_6_-EsAgo WT and (**D**) His_6_-EsAgo DM were purified by FPLC following 5 hours of IPTG induction. Cell lysate was used as the input (In) for the HisTrap (HT) IMAC column. Eluate fractions from the HT column with His_6_-EsAgo also contained proteins of various sizes and were therefore pooled, concentrated, and used as the input for the SEC column. Proteins purified via SEC were analyzed on 12% SDS-PAGE gels stained with Coomassie. The thick ~75 kDa bands correspond to purified WT and DM His_6_-EsAgo. The cleanest fractions are highlighted in red and were subsequently used for *in vitro* assays. The largest peaks in UV absorbance at 280 nm displayed on the chromatograms correspond to fractions containing the proteins of interest.

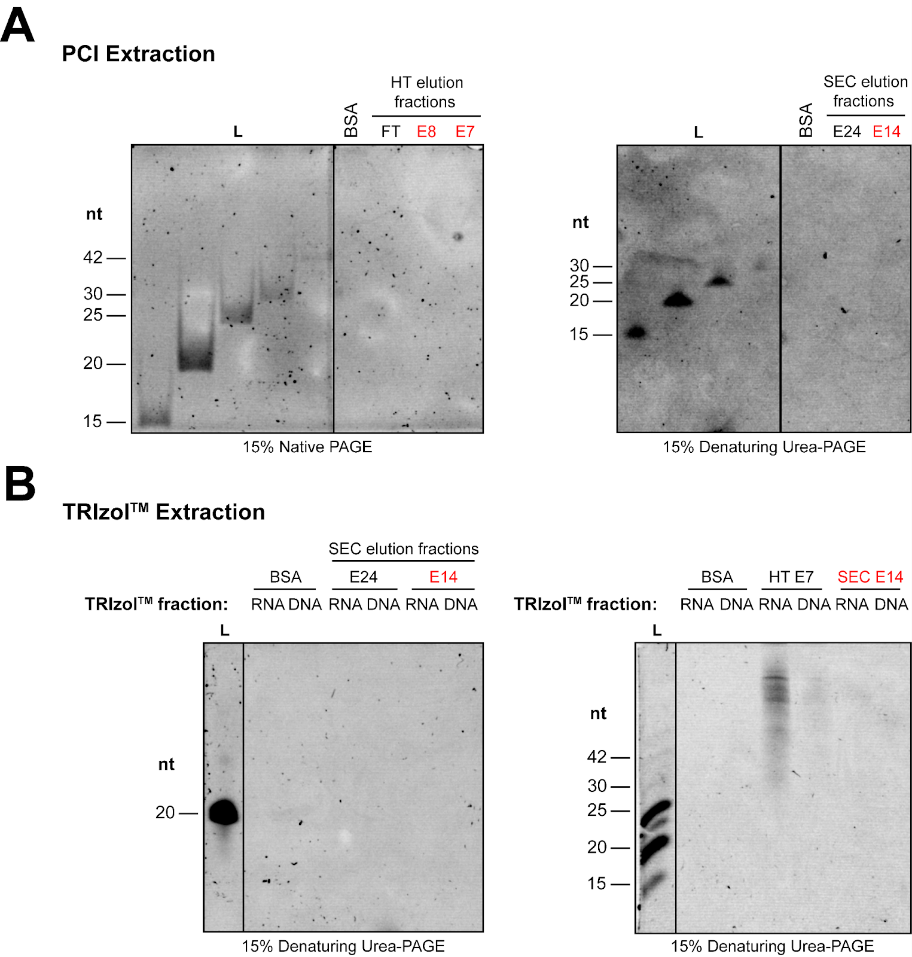

**Supplementary Fig. 6. His_6_-EsAgo WT may purify free of associated short nucleic acid guides following overexpression in *E. coli***. 500 pmols of His_6_-EsAgo WT was digested with proteinase K and any nucleic acids present were extracted by way of (**A**) PCI or (**B**) TRIzol^TM^ precipitation. PCI extracts represent total nucleic acid fractions whereas extraction with TRIzol^TM^ Reagent separates nucleic acids into DNA and RNA fractions. Extracted nucleic acids were resolved on 15% native PAGE or 15% denaturing PAGE gels with 7 M urea. Fractions containing His_6_-EsAgo WT are highlighted in red. An equivalent amount of BSA and proteins from other fractions, such as flow through (FT) eluted from the HT column, served as negative controls. As pAgo guides are short (~15-30 nt) and single-stranded, primers of known length were used as molecular weight markers.

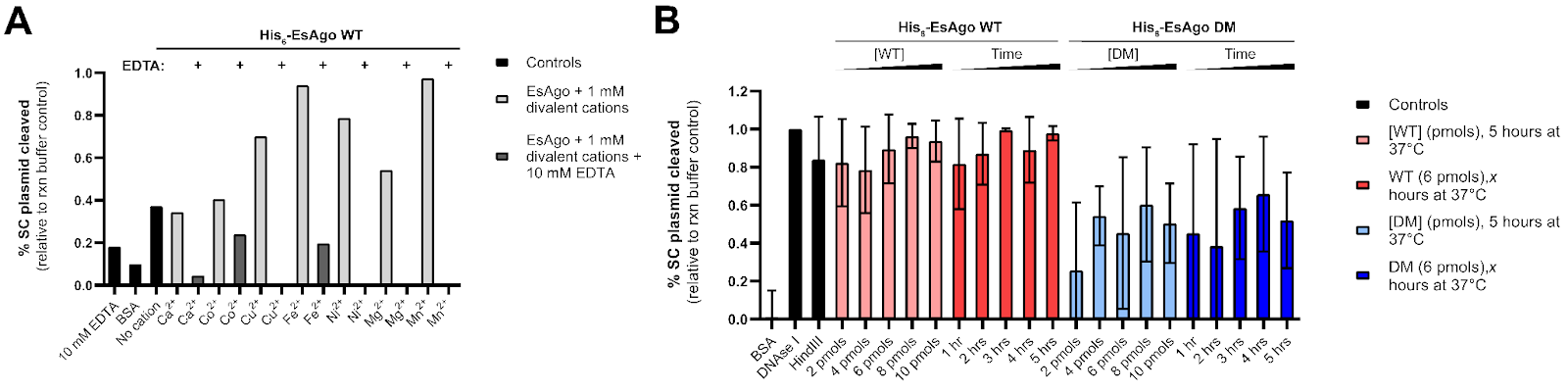

**Supplementary Fig. 7. Plasmids cleaved from unguided EsAgo assays.** (**A**) Percent supercoiled (SC) plasmid cleaved when His_6_-EsAgo was supplied with 1 mM of various divalent cations (refer to **Fig. 5A**). Values were determined relative to the intensity of the supercoiled band of pQE-80L incubated in reaction (rxn) buffer only. Band intensities were measured using ImageJ. (**B**) Percent supercoiled (SC) plasmid cleaved relative to the supercoiled band of pQE-80L incubated in reaction buffer only (refer to **Fig. 5B**). Error bars indicate SD of three technical replicates per reaction (n=3).

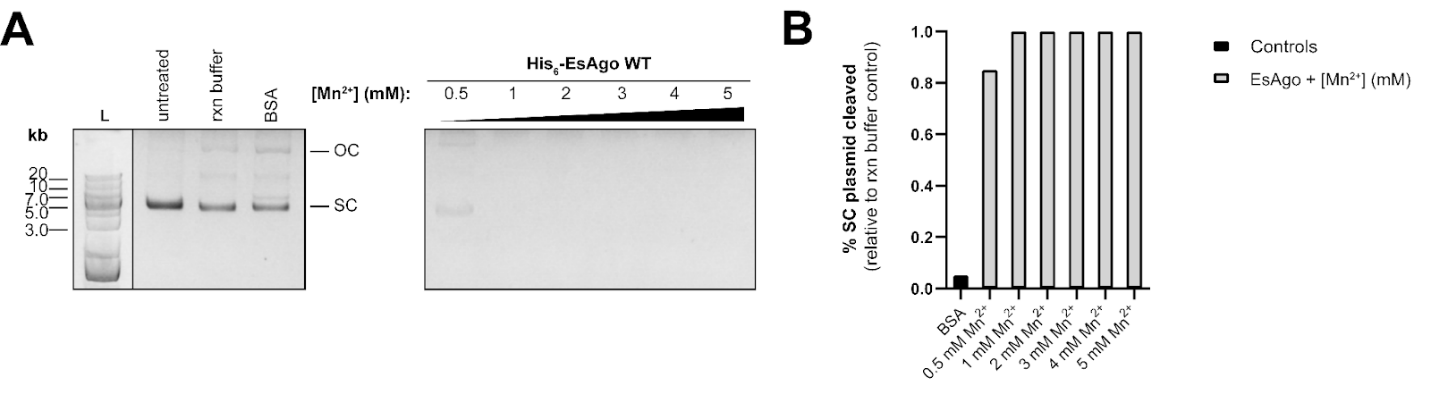

**Supplementary Fig. 8. EsAgo efficiently degrades plasmids *in vitro* with Mn^2+^**. (**A**) Effects of different concentrations of Mn^2+^ on His_6_-EsAgo-mediated DNA cleavage. Reactions were incubated at 37°C for 5 hours at a EsAgo:pQE-80L ratio of 1 pmol:12.5 ng then resolved on an AGE gel. (**B**) Percent supercoiled plasmid cleaved when EsAgo is supplied with various concentrations of Mn^2+^ (n=1). Values were determined relative to the supercoiled band of pQE-80L incubated in reaction buffer only.

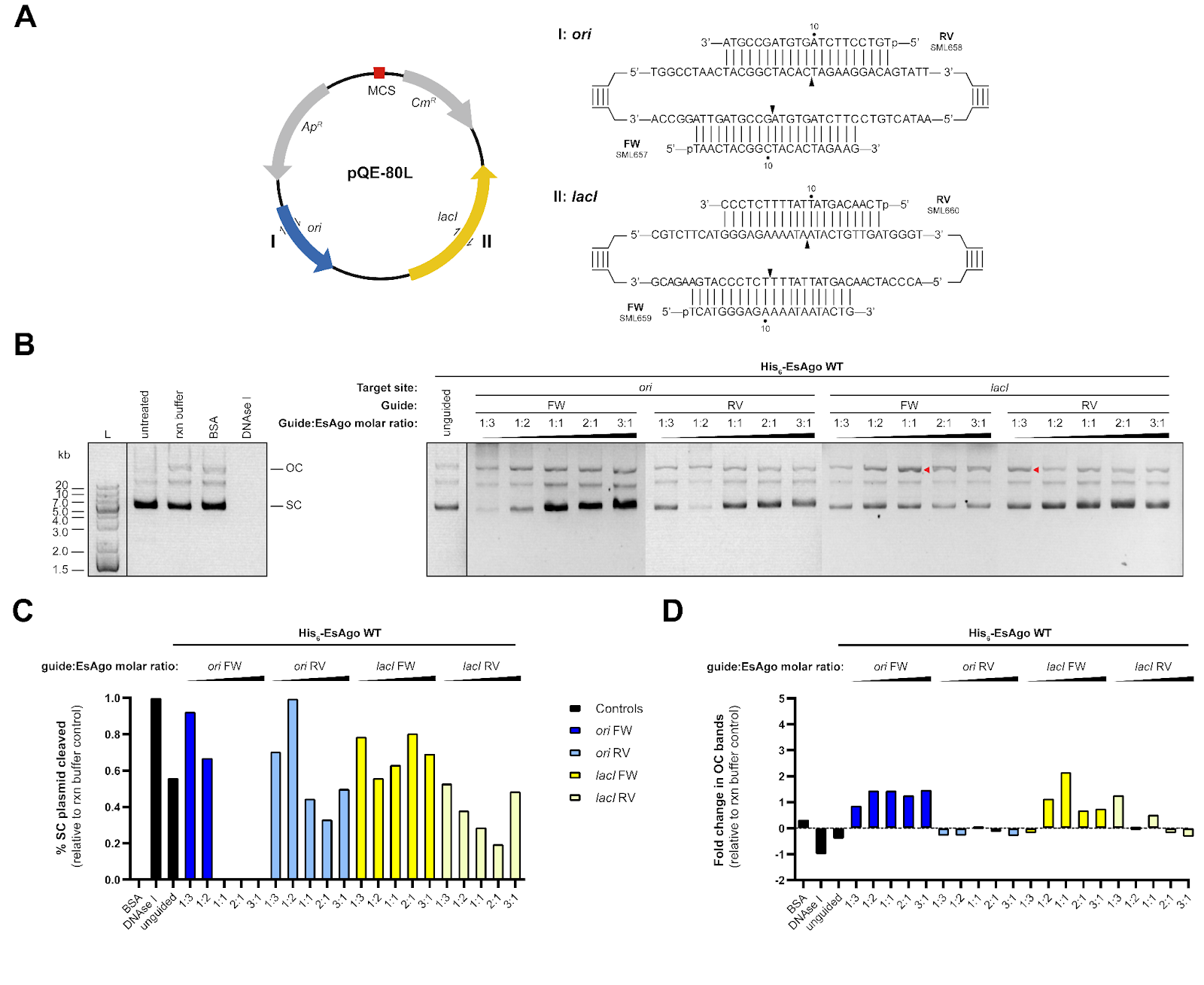

**Supplementary Fig. 9. Efficiency of programmed cleavage may depend on an appropriate guide:EsAgo molar ratio.** (**A**) Guides consisted of 21 nt-long ssDNA oligos with 5’-phosphate groups targeting regions with lower GC content in the *ori* (44% GC) and *lacI* (32% GC) genes of pQE-80L. Black triangles indicate the expected cleavage site of each guide. (**B**) Dose response analysis when providing His_6_-EsAgo WT with individual forward or reverse guides to test for guide functionality. Single-stranded cleavage from an EsAgo-guide complex was expected to convert supercoiled plasmids into an open-circular (OC) state in a dose-dependent manner. Reactions consisted of 500 ng of plasmid incubated for 5 hours at 37°C with 30 pmols of EsAgo WT and various amounts of guides. Plasmids incubated with 30 pmols of BSA and DNase I served as negative and positive controls, respectively, for DNA degradation. Red arrows indicate notable instances of an increase in open-circular plasmids. (**C**) Percent supercoiled (SC) pQE-80L cleaved and (**D**) fold change of pQE-80L in the open-circular state determined relative to the supercoiled and open-circular band, respectively, of pQE-80L incubated in reaction buffer only (n=1). Guides targeting *ori* were not used for subsequent dsDNA cleavage assays as the *ori*-RV guide appeared nonfunctional. Note: Despite exact replicates of this assay, increases in the concentration of open-circular bands (with an appropriate guide:EsAgo molar ratio) and attenuation of random plasmid degradation were repeatedly observed during other guided assays.

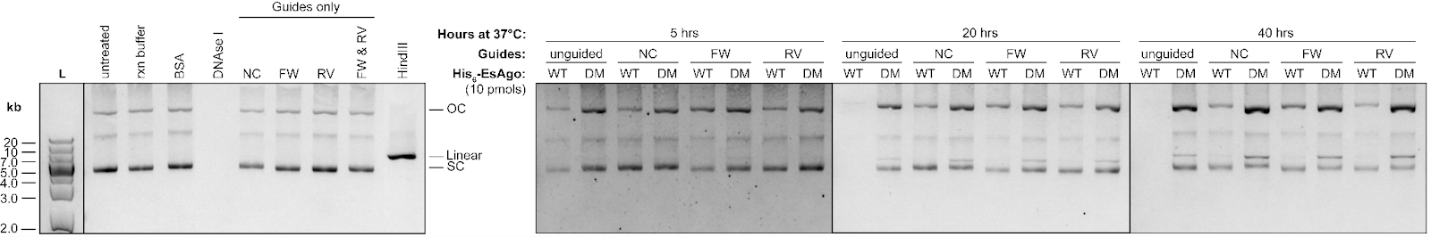

**Supplementary Fig. 10. Supplying EsAgo with one 5’-P ssDNA guide attenuates nonspecific plasmid degradation**. Reactions consisted of 10 pmols of His_6_-EsAgo WT- and DM-guide complexes incubated with 100 ng of pQE-80L for 5, 20, and 40 hours. The guide pair targeting pQE-80L *lacI* was used for this assay. Forward and non-complementary (NC) guides were supplied to EsAgo WT and DM at a molar ratio of 1:1 while reverse guides were supplied at a molar ratio of 1 guide:3 EsAgo. Control reactions depicted on the AGE gel on the left are from the 40-hour incubation setup.

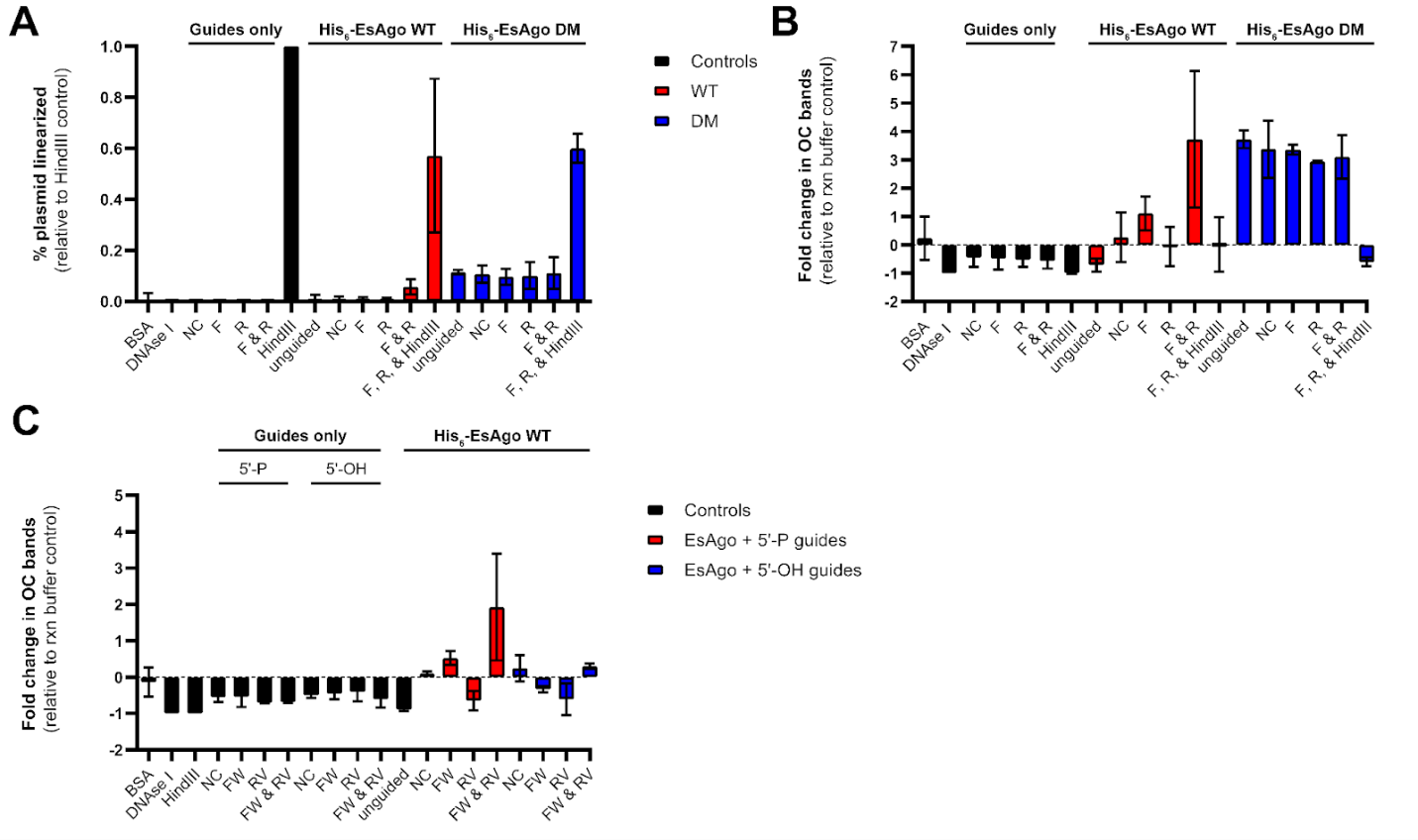

**Supplementary Fig. 11. Plasmids cleaved from guided EsAgo assays.** (**A**) Percent plasmid linearized and (**B**) fold change of open-circular (OC) pQE-80L relative to HindIII digests (i.e. 100% plasmids linearization) and the open-circular band of pQE-80L incubated in reaction buffer only, respectively, when His_6_-EsAgo WT and DM were supplied with 5’-P *lacI* guides (refer to **Fig. 6B**). Scrambled guides served as non-complementary (NC) controls. Error bars indicate SD of technical replicates (n=4 for controls and EsAgo WT) (n=2 for EsAgo DM). (**C**) fold change in open-circular plasmid bands relative to the open-circular band of pQE-80L incubated in reaction buffer only when EsAgo was supplied with 5’-P or 5’-OH *lacI* guides (refer to **Fig. 6C**). Error bars indicate SD of technical replicates (n=2).

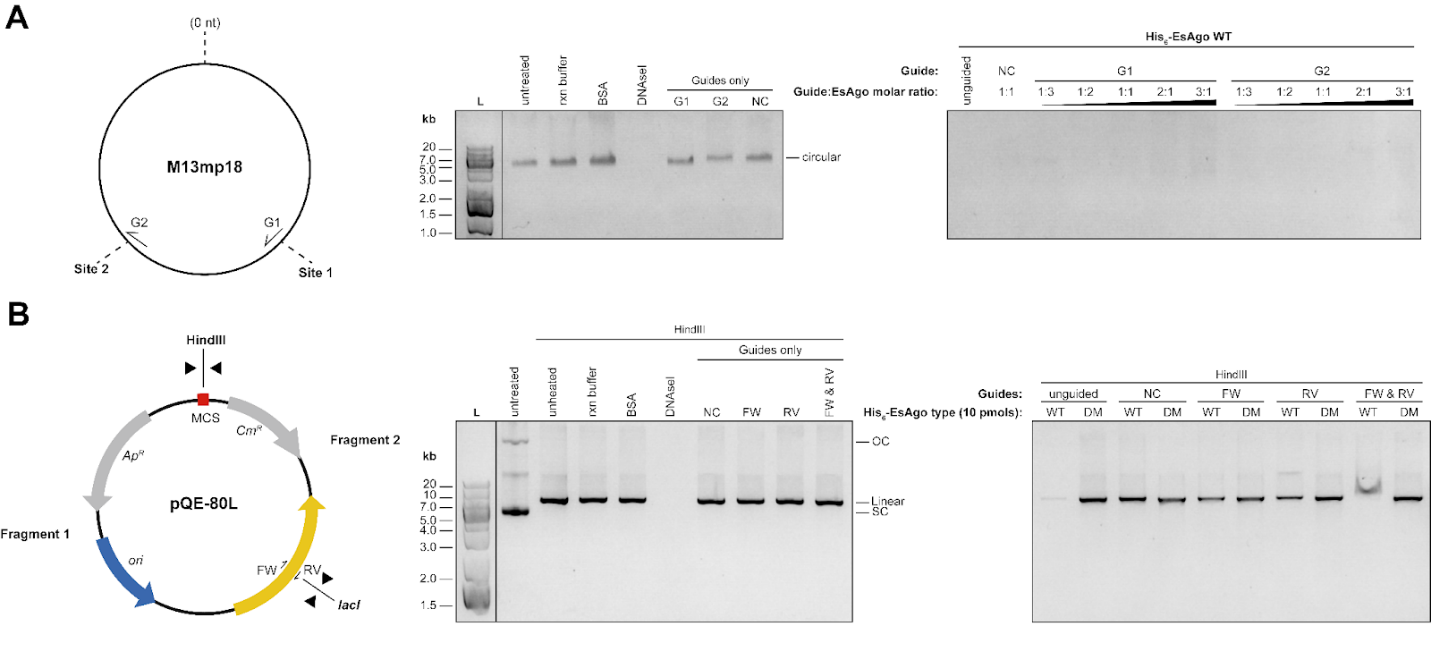

**Supplementary Fig. 12. EsAgo degrades circular ssDNA and linear dsDNA substrates in a random guide-independent manner.** (**A**) Guided cleavage assay of ssDNA targeting 2 different sites on the M13mp18 phage genome. Reactions consisted of 50 ng of M13mp18 incubated with 5 pmols of His_6_-EsAgo WT for 5 hours at 37°C. Guides were loaded onto EsAgo at various guide:EsAgo molar ratios. Phage ssDNA was randomly degraded by EsAgo-guide complexes across all reactions and in subsequent attempts at targeting M13mp18 for cleavage (not shown). (**B**) Guided cleavage assay targeting the *lacI* site of HindIII-linearized pQE-80L. Reactions consisted of 100 ng of pre-linearized plasmids incubated with 10 pmols of His_6_-EsAgo WT or DM. Guides were supplied at guide:EsAgo molar ratios of 1:1 for NC and FW guides, and 1:3 for RV guides. Expected fragments ~1.6 kb and ~3.1 kb in size following dsDNA cleavage at the intended site were not detected. Plasmids and phage ssDNA incubated with BSA or DNAseI served as negative and positive controls, respectively, for nuclease-mediated DNA degradation.

**Supplementary Table 1. Contigs containing pAgos from various *Exiguobacterium* species used for multiple genome alignment.**

| **Host *Exiguobacterium* species** | **Contig NCBI accession** | **pAgo location on contig** | **pAgo NCBI accession** | **pAgo % identity with EsAgo** | **Remarks** | **Contig Source** |
| --- | --- | --- | --- | --- | --- | --- |
| sp. AB2* | JNAA01000033.1 | 33,959-36,067 (+) | WP_034806158.1 |  | EsAgo | [2] |
| sp. s150* | JACSMC010000014.1 | 51,801-53,909 (+) | WP_214844161.1 | 96.15% |  | [57] |
| sp. IPCH1* | SJVE01000013.1 | 51,758-53,866 (+) | WP_131472818.1 | 95.73% | Identical pAgos are encoded on these contigs | Castro-Severyn et al. (NCBI database) |
| sp. IPCI3* | SJVD01000013.1 | 51,695-53,803 (+) |  |  |  |  |
| sp. IPBC4* | SJVF01000013.1 | 51,758-53,866 (+) |  |  |  |  |
| sp. s157* | JACSLQ010000008.1 | 41,047-43,155 (+) | WP_214756792.1 | 94.87% |  | [57] |
| *aurantiacum** | JAHXHC010000004.1 | 72,024-74,129 (-) | WP_233005385.1 | 93.30% |  | [58] |
| *qingdaonense** | JACSKY010000007.1 | 72,464-74,569 (-) | WP_215144538.1 | 88.46% |  | [57] |
| sp. s26* | JACSLS010000010.1 | 18,217-20,325 (+) | WP_214828248.1 | 94.16% |  |  |
| sp. MER 193* | JAMAVV010000021.1 | 16,578-18,686 (+) | WP_251307042.1 | 89.46% |  | [59] |
| sp. s163* | JACSJN010000008.1 | 63,997-66,102 (-) | WP_214689493.1 | 89.03% |  | [57] |
| sp. s21* | JACSLF010000009.1 | 60,737-62,845 (+) | WP_214788950.1 | 95.73% |  |  |
| sp. SH3S1* | SJUW01000007.1 | 98,954-101,062 (-) | WP_161567616.1 | 93.45% |  | Castro-Severyn et al. (NCBI database) |
| sp. s55* | JACSLE010000011.1 | 60,030-62,138 (+) | WP_214712037.1 | 95.44% |  | [57] |
| sp. CinTr1* | CANDNX010000009.1 | 62,648-64,756 (-) | WP_270813360.1 | 89.60% |  | Costa & Keller-Costa (NCBI database) |
| sp. s7 | JACSLO010000010.1 | 55,157-57,265 (-) | WP_215192561.1 | 95.30% |  | [57] |
| sp. AT1b | MOEL01000004.1 | 53,075-55,183 (-) | WP_074032557.1 | 89.32% |  | [60] |
| sp. JLM-2 | LDNY01000004.1 | 78,280-80,097 (-) | WP_015880612.1 | 88.46% |  | [61] |
| sp. N5 | JAPAEP010000004.1 | 78,745-80,853 (-) | WP_264093295.1 | 89.70% |  | [62] |
| sp. s48 | JACSKC010000004.1 | 79,247-81,352 (-) | WP_214742983.1 | 88.89% |  | [57] |
| *aestuarii* | JAIVAC010000004.1 | 36,146-38,251 (+) | WP_224863494.1 | 87.89%^***^ |  | Yu et al. (NCBI database) |
| *marinum* | JHZT01000001.1 | 92,041-94,146 (-) | WP_026824436.1 | 89.46% | Contig contains EmaAgo [18] | [63] |
| sp. s38 | JACSKT010000005.1 | 84,234-85,484 (-) | WP_214704126.1 | 91.77% | Identical pAgos are encoded on these contigs | [57] |
| sp. s89 | JACSKS010000005.1 | 84,234-85,484 (-) |  |  |  |  |
| sp. s139 | JACSKU010000005.1 | 29,658-30,908 (+) |  |  |  |  |
| sp. s196 | JACSJR010000010.1 | 33,289-35,397 (+) | WP_214814469.1 | 92.31% |  |  |
| sp. s149 | JACSLZ010000009.1 | 24,185-26,293 (+) | WP_214720089.1 | 95.16% |  |  |
| sp. SL-10 | SJUP01000008.1 | 56,822-58,930 (+) | WP_131436798.1 | 97.01%^**^ |  | Castro-Severyn et al. (NCBI database) |
| sp. s181 | JACSJM010000005.1 | 63,916-66,024 (-) | WP_214812784.1 | 92.17% |  | [57] |
| sp. s140 | JACSJK010000006.1 | 64,567-66,672 (-) | WP_214873908.1 | 89.17% |  |  |

Contigs are ordered based on their appearance in **Supplementary Fig. 1**.  “*” The general organization of the contigs derived from these species is similar to that of the contig containing *EsAgo* and are displayed in **Fig. 2B**. Sequence similarity of the various *Exiguobacterium* pAgos to EsAgo are listed wherein “**” and “***” indicate the highest and lowest percent identities observed, respectively. Note: Contigs are identical between *E.* sp. IPCH1, IPCI3, and IPBC4, and between sp. s38, s89, and s139.

**Supplementary Table 2. Putative defense systems identified by PADLOC on *Exiguobacterium* contigs.**

| **Host *Exiguobacterium* species** | **Putative defense systems on contig** | | | | | | | | |
| --- | --- | --- | --- | --- | --- | --- | --- | --- | --- |
|  | **system** | **protein.name** | **target.name** | **target.description** | **seqid** | **start** | **end** | **strand** | **relative.position** |
| **sp. AB2*** | wadjet_other | JetA | DI14_09635 | DI14_09635 | JNAA01000033.1 | 2027 | 3503 | + | 4 |
|  | wadjet_other | JetC | DI14_09645 | DI14_09645 | JNAA01000033.1 | 4674 | 8745 | + | 5 |
|  | wadjet_other | JetD | DI14_09650 | DI14_09650 | JNAA01000033.1 | 8741 | 10019 | + | 6 |
|  | RM_type_l | MTase_l | DI14_09670 | DI14_09670 | JNAA01000033.1 | 11629 | 13642 | + | 9 |
|  | RM_type_l | REase_l | DI14_09685 | DI14_09685 | JNAA01000033.1 | 14910 | 17901 | + | 12 |
|  | DMS_other | DrmB | DI14_09690 | DI14_09690 | JNAA01000033.1 | 17969 | 19067 | - | 13 |
|  | DMS_other | DrmA | DI14_09700 | DI14_09700 | JNAA01000033.1 | 19710 | 23121 | - | 14 |
|  | argonaute_solo | pAgo | DI14_09730 | DI14_09730 | JNAA01000033.1 | 33958 | 36067 | + | 20 |
| **sp. s150*** | wadjet_type_ll | JetA2 | JACSMC010000014.1_32 | 22534 24009 1 ID=1_32;partial=00;start_type=ATG;rbs_motif=GGAG/GAGG;rbs_spacer=5-10bp;gc_cont=0.520 | JACSMC010000014.1 | 22534 | 24009 | + | 32 |
|  | wadjet_type_ll | JetB2 | JACSMC010000014.1_33 | 24006 25160 1 ID=1_33;partial=00;start_type=ATG;rbs_motif=AGGAG;rbs_spacer=5- 10bp;gc_cont=0.516 | JACSMC010000014.1 | 24006 | 25160 | + | 33 |
|  | wadjet_type_ll | JetC2 | JACSMC010000014.1_34 | 25157 29251 1 ID=1_34;partial=00;start_type=ATG;rbs_motif=AGGAG;rbs_spacer=5-10bp;gc_cont=0.536 | JACSMC010000014.1 | 25157 | 29251 | + | 34 |
|  | wadjet_type_ll | JetD2 | JACSMC010000014.1_35 | 29248 30528 1 ID=1_35;partial=00;start_type=ATG;rbs_motif=GGA/GAG/AGG;rbs_spacer=5-10bp;gc_cont=0.533 | JACSMC010000014.1 | 29248 | 30528 | + | 35 |
|  | RM_type_l | MTase_l | JACSMC010000014.1_38 | 32135 34147 1 ID=1_38;partial=00;start_type=ATG;rbs_motif=GGAG/GAGG;rbs_spacer=5-10bp;gc_cont=0.386 | JACSMC010000014.1 | 32135 | 34147 | + | 38 |
|  | RM_type_I | Specificity_l | JACSMC010000014.1_39 | 34144 35439 1 ID=1_39:partial=00;start_type=ATG;rbs_motif=AGxAGG/AGGxGG;rbs_spacer=5- 10bp;gc_cont=0.332 | JACSMC010000014.1 | 34144 | 35439 | + | 39 |
|  | RM_type_I | REaseJ | JACSMC010000014.1_40 | 35454 38444 1 ID=1_40:partial=00;start_type=ATG;rbs_motif=GGAG/GAGG;rbs_spacer=5-10bp;gc_cont=0.416 | JACSMC010000014.1 | 35454 | 38444 | + | 40 |
|  | DMS_other | DrmB | JACSMC010000014.1_41 | 38510 40264-1 ID=1_41;partial=00;start_type=ATG;rbs_motif=AGGAG;rbs_spacer=5-10bp;gc_cont=0.425 | JACSMC010000014.1 | 38510 | 40264 | - | 41 |
|  | DMS_other | DrmA | JACSMC010000014.1_42 | 40251 43661 -1 ID=1_42;partial=00;start_type=ATG;rbs_motif=GGA/GAG/AGG;rbs_spacer=5-10bp;gc_cont=0.429 | JACSMC010000014.1 | 40251 | 43661 | - | 42 |
|  | argonaute_solo | pAgo | JACSMC010000014.1_47 | 51801 53909 1 ID=1_47;partial=00;start_type=ATG;rbs_motif=AGGA;rbs_spacer=5- 10bp;gc_cont=0.476 | JACSMC010000014.1 | 51801 | 53909 | + | 47 |
| **sp. IPCH1*** | wadjet_type_ll | JetA2 | EVJ18_12810 | EVJ18_12810 | SJVE01000013.1 | 25256 | 26732 | + | 31 |
|  | wadjet_type_ll | JetB2 | EVJ 1812815 | EVJ18_12815 | SJVE01000013.1 | 26728 | 27883 | + | 32 |
|  | wadjet_type_ll | JetC2 | EVJ18_12820 | EVJ18_12820 | SJVE01000013.1 | 27771 | 31974 | + | 33 |
|  | wadjet_type_ll | JetD2 | EVJ18_12825 | EVJ 18_12825 | SJVE01000013.1 | 31970 | 33251 | + | 34 |
|  | RM_type_l | MTase_l | EVJ18_12840 | EVJ 18_12840 | SJVE01000013.1 | 34857 | 36870 | + | 37 |
|  | RM_type_l | Specificity_l | EVJ18_12845 | EVJ 18_12845 | SJVE01000013.1 | 36866 | 38192 | + | 38 |
|  | RM_type_l | REase_l | EVJ 18_12850 | EVJ 18_12850 | SJVE01000013.1 | 38206 | 41197 | + | 39 |
|  | DMS_other | DrmB | EVJ 18_12855 | EVJ 18_12855 | SJVE01000013.1 | 41265 | 43020 | - | 40 |
|  | DMS_other | DrmA | EVJ18_12860 | EVJ 18_12860 | SJVE01000013.1 | 43006 | 46417 | - | 41 |
|  | argonaute_solo | pAgo | EVJ18 12885 | EVJ 18_12885 | SJVE01000013.1 | 51757 | 53866 | + | 46 |
| **sp. IPCI3*** | wadjet_type_ll | JetA2 | EVJ19_12820 | EVJ19_12820 | SJVD01000013.1 | 25193 | 26669 | + | 31 |
|  | wadjet_type_ll | JetB2 | EVJ 19_12825 | EVJ 19_12825 | SJVD01000013.1 | 26665 | 27820 | + | 32 |
|  | wadjet_type_ll | JetC2 | EVJ 19_12830 | EVJ 19_12830 | SJVD01000013.1 | 27708 | 31911 | + | 33 |
|  | wadjet_type_ll | JetD2 | EVJ 19_12835 | EVJ19_12835 | SJVD01000013.1 | 31907 | 33188 | + | 34 |
|  | RM_type_l | MTase_l | EVJ 19_12850 | EVJ 19_12850 | SJVD01000013.1 | 34794 | 36807 | + | 37 |
|  | RM_type_l | SpecificityJ | EVJ 19_12855 | EVJ 19_12855 | SJVD01000013.1 | 36803 | 38129 | + | 38 |
|  | RM_type_l | REase_l | EVJ 19_12860 | EVJ 19_12860 | SJVD01000013.1 | 38143 | 41134 | + | 39 |
|  | DMS_other | DrmB | EVJ 19_12865 | EVJ19_12865 | SJVD01000013.1 | 41202 | 42957 | - | 40 |
|  | DMS_other | DrmA | EVJ 19_12870 | EVJ 19_12870 | SJVD01000013.1 | 42943 | 46354 | - | 41 |
|  | argonaute_solo | pAgo | EVJ 19_12895 | EVJ 19_12895 | SJVD01000013.1 | 51694 | 53803 | + | 46 |
| **sp. IPBC4*** | wadjet_type_ll | JetA2 | EVJ17_12810 | EVJ17_12810 | SJVF01000013.1 | 25256 | 26732 | + | 31 |
|  | wadjet_type_ll | JetB2 | EVJ17_12815 | EVJ 1712815 | SJVF01000013.1 | 26728 | 27883 | + | 32 |
|  | wadjet_type_ll | JetC2 | EVJ17_12820 | EVJ17_12820 | SJVF01000013.1 | 27771 | 31974 | + | 33 |
|  | wadjet_type_ll | JetD2 | EVJ 17_12825 | EVJ 17_12825 | SJVF01000013.1 | 31970 | 33251 | + | 34 |
|  | RM_type_l | MTase_l | EVJ 17_12840 | EVJ 17_12840 | SJVF01000013.1 | 34857 | 36870 | + | 37 |
|  | RM_type_l | SpecificityJ | EVJ 17_12845 | EVJ 17_12845 | SJVF01000013.1 | 36866 | 38192 | + | 38 |
|  | RM_type_l | REaseJ | EVJ 17_12850 | EVJ17_12850 | SJVF01000013.1 | 38206 | 41197 | + | 39 |
|  | DMS_other | DrmB | EVJ 17_12855 | EVJ 17_12855 | SJVF01000013.1 | 41265 | 43020 | - | 40 |
|  | DMS_other | DrmA | EVJ 17_12860 | EVJ 17_12860 | SJVF01000013.1 | 43006 | 46417 | - | 41 |
|  | argonaute_solo | pAgo | EVJ 17_12885 | EVJ 17_12885 | SJVF01000013.1 | 51757 | 53866 | + | 46 |
| **sp. s157*** | RM_type_l | MTase_l | JACSLQ010000008.1_34 | 24146 26158 1 ID=1_34;partial=00;start_type=ATG;rbs_motif=GGAG/GAGG;rbs_spacer=5-10bp;gc_cont=0.380 | J ACSLQ010000008.1 | 24146 | 26158 | + | 34 |
|  | RM_type_l | Specificity_l | J ACSLQ010000008.1 _35 | 26155 27480 1 ID=1_35;partial=00;start_type=ATG;rbs_motif=GGAG/GAGG;rbs_spacer=5- 10bp;gc_cont=0.314 | J ACSLQ010000008.1 | 26155 | 27480 | + | 35 |
|  | RM_type_l | REasel | JACSLQ010000008.1_36 | 27495 30485 1 ID=1_36;partial=00;start_type=ATG;rbs_motif=GGAG/GAGG;rbs_spacer=5- 10bp;gc_cont=0.419 | JACSLQ010000008.1 | 27495 | 30485 | + | 36 |
|  | DMS_other | DrmB | JACSLQ010000008.1_37 | 30554 32308 -1 ID=1_37;partial=00;start_type=ATG;rbs_motif=AGGAG;rbs_spacer=5- 10bp;gc_cont=0.421 | JACSLQ010000008.1 | 30554 | 32308 | - | 37 |
|  | DMS_other | DrmA | JACSLQ010000008.1_38 | 32295 35705 -1 ID=1_38;partial=00;start_type=ATG;rbs_motif=GGA/GAG/AGG;rbs_spacer=5-10bp;gc_cont=0.433 | J ACSLQ010000008.1 | 32295 | 35705 | - | 38 |
|  | argonaute_solo | pAgo | JACSLQ010000008.1_43 | 41047 43155 1 ID=1_43;partial=00;start_type=ATG;rbs_motif=GGAG/GAGG;rbs_spacer=5-10bp;gc_cont=0.459 | J ACSLQ010000008.1 | 41047 | 43155 | + | 43 |
| ***aurantiacum****** | argonaute_solo | pAgo | JAHXHC010000004.1_85 | 72024 74129-1 ID=1_85;partial=00;start_type=ATG;rbs_motif=GGAG/GAGG;rbs_spacer=5- 10bp;gc_cont=0.460 | JAHXHC010000004.1 | 72024 | 74129 | - | 85 |
|  | DMS_other | DrmA | JAHXHC010000004.1_90 | 79471 82881 1 ID=1_90;partial=00;start_type=ATG;rbs_motif=GGA/GAG/AGG;rbs_spacer=5- 10bp;gc_cont=0.431 | JAHXHC010000004.1 | 79471 | 82881 | + | 90 |
|  | DMS_other | DrmB | JAHXHC010000004.1_91 | 82868 84622 1 ID=1_91;partial=00;start_type=ATG;rbs_motif=AGGAG;rbs_spacer=5- 10bp;gc_cont=0.423 | JAHXHC010000004.1 | 82868 | 84622 | + | 91 |
|  | RM_type_l | REase_l | JAHXHC010000004.1_92 | 84691 87681 -1 ID=1_92;partial=00;start_type=ATG;rbs_motif=GGAG/GAGG;rbs_spacer=5- 10bp;gc_cont=0.417 | JAHXHC010000004.1 | 84691 | 87681 | - | 92 |
|  | RMJypeJ | SpecificityJ | JAHXHC010000004.1_93 | 87697 89001 -1 ID=1_93;partial=00;start_type=ATG;rbs_motif=AGxAGG/AGGxGG;rbs_spacer=5- 10bp;gc_cont=0.294 | JAHXHC010000004.1 | 87697 | 89001 | - | 93 |
|  | RM_type_l | MTase_l | JAHXHC010000004.1 _94 | 88998 91010-1 ID=1_94;partial=00;start_type=ATG;rbs_motif=GGAG/GAGG;rbs_spacer=5- 10bp;gc_cont=0.377 | JAHXHC010000004.1 | 88998 | 91010 | - | 94 |
| ***qingdaonense**** | cbass_type_l | Cyclase | J ACSKY010000007.1 _7 | 8587 9957 -1 ID=1_7;partial=00;start_type=TTG;rbs_motif=GGAG/GAGG;rbs_spacer=5-10bp;gc_cont=0.334 | JACSKY010000007.1 | 8587 | 9957 | - | 7 |
|  | cbass_type_l | Effector | J ACSKY010000007.1_8 | 9972 11075 -1 ID=1_8;partial=00;start_type=ATG;rbs_motif=GGxGG;rbs_spacer=5-10bp;gc_cont=0.308 | JACSKY010000007.1 | 9972 | 11075 | - | 8 |
|  | cas_type_l-C | Cas2c | J ACSKY010000007.1_37 | 37100 37390 -1 ID=1_37;partial=00;start_type=ATG;rbs_motif=GGAG/GAGG;rbs_spacer=5-10bp;gc_cont=0.399 | JACSKY010000007.1 | 37100 | 37390 | - | 37 |
|  | cas_type_l-C | Caslc | JACSKY010000007.1_38 | 37401 38432 -1 ID=1_38;partial=00;start_type=TTG;rbs_motif=AGGAG;rbs_spacer=5-10bp;gc_cont=0.409 | JACSKY010000007.1 | 37401 | 38432 | - | 38 |
|  | cas_type_l-C | Cas4c | JACSKY010000007.1_39 | 38429 39088 -1 ID=1_39;partial=00;start_type=ATG;rbs_motif=None;rbs_spacer=None;gc_cont=0.409 | JACSKY010000007.1 | 38429 | 39088 | - | 39 |
|  | cas_type_l-C | Cas7c | J ACSKY010000007.1 _40 | 39078 39932 -1 ID=1_40;partial=00;start_type=ATG;rbs_motif=AGGAG/GGAGG;rbs_spacer=11-12bp;gc_cont=0.428 | JACSKY010000007.1 | 39078 | 39932 | - | 40 |
|  | cas_type_l-C | Cas8c | JACSKY010000007.1_41 | 39929 41806 -1 ID=1_41;partial=00;start_type=ATG;rbs_motif=AGxAGG/AGGxGG;rbs_spacer=5-10bp;gc_cont=0.407 | JACSKY010000007.1 | 39929 | 41806 | - | 41 |
|  | cas_type_l-C | Cas5c | J ACSKY010000007.1_42 | 41818 42516 -1 ID=1_42;partial=00;start_type=ATG;rbs_motif=AGGAG;rbs_spacer=5-10bp;gc_cont=0.416 | JACSKY010000007.1 | 41818 | 42516 | - | 42 |
|  | cas_type_l-C | Cas3c | JACSKY010000007.1_43 | 42533 44866 -1 ID=1_43;partial=00;start_type=GTG;rbs_motif=GGA/GAG/AGG;rbs_spacer=5-10bp;gc_cont=0.404 | JACSKY010000007.1 | 42533 | 44866 | - | 43 |
|  | argonaute_solo | pAgo | J ACSKY010000007.1_80 | 72464 74569 -1 ID=1_80;partial=00;start_type=ATG;rbs_motif=GGAG/GAGG;rbs_spacer=5-10bp;gc_cont=0.445 | JACSKY010000007.1 | 72464 | 74569 | - | 80 |
|  | DMS_other | DrmA | J ACSKY010000007.1_86 | 82187 85597 1 ID=1_86;partial=00;start_type=ATG;rbs_motif=GGA/GAG/AGG;rbs_spacer=5-10bp;gc_cont=0.413 | JACSKY010000007.1 | 82187 | 85597 | + | 86 |
|  | DMS_other | DrmB | J ACSKY010000007.1 _87 | 85584 87338 1 ID=1_87;partial=00;start_type=ATG;rbs_motif=AGGAG;rbs_spacer=5-10bp;gc_cont=0.423 | JACSKY010000007.1 | 85584 | 87338 | + | 87 |
|  | RM_type_l | REase_l | JACSKY010000007.1_88 | 87408 90398 -1 ID=1_88;partial=00;start_type=ATG;rbs_motif=GGAG/GAGG;rbs_spacer=5-10bp;gc_cont=0.414 | JACSKY010000007.1 | 87408 | 90398 | - | 88 |
|  | RM_type_l | SpecificityJ | JACSKY010000007.1_89 | 90413 91717 -1 ID=1_89;partial=00;start_type=ATG;rbs_motif=AGxAGG/AGGxGG;rbs_spacer=5-10bp;gc_cont=0.290 | JACSKY010000007.1 | 90413 | 91717 | - | 89 |
|  | RM_type_l | MTaseJ | JACSKY010000007.1_90 | 91714 93726 -1 ID=1_90;partial=00;start_type=ATG;rbs_motif=GGAG/GAGG;rbs_spacer=5-10bp;gc_cont=0.373 | JACSKY010000007.1 | 91714 | 93726 | - | 90 |
|  | wadjet_type_ll | JetD2 | JACSKY010000007.1_93 | 95337 96611 -1 ID=1_93;partial=00;start_type=ATG;rbs_motif=GGA/GAG/AGG;rbs_spacer=5-10bp;gc_cont=0.482 | JACSKY010000007.1 | 95337 | 96611 | - | 93 |
|  | wadjet_type_ll | JetC2 | J ACSKY010000007.1_94 | 96608 100702 -1 ID=1_94;partial=00;start_type=ATG;rbs_motif=AGGAG/GGAGG;rbs_spacer=11-12bp;gc_cont=0.459 | JACSKY010000007.1 | 96608 | 1E+05 | - | 94 |
|  | wadjet_type_ll | JetB2 | J ACSKY010000007.1_95 | 100699 101856 -1 ID=1_95;partial=00;start_type=ATG;rbs_motif=AGGAG;rbs_spacer=5-10bp;gc_cont=0.441 | JACSKY010000007.1 | 1E+05 | 1E+05 | - | 95 |
|  | wadjet_type_ll | JetA2 | J ACSKY010000007.1_96 | 101853 103328 -1 ID=1_96;partial=00;start_type=ATG;rbs_motif=AGGAG;rbs_spacer=5-10bp;gc_cont=0.434 | JACSKY010000007.1 | 1E+05 | 1E+05 | - | 96 |
| **sp. s26*** | DMS_other | Specificity_l | JACSLS010000010.1 2 | 1063 2385 1 ID=1_2;partial=00;start_type=ATG;rbs_motif=AGxAGG/AGGxGG;rbs_spacer=5-10bp;gc_cont=0.308 | JACSLS010000010.1 | 1063 | 2385 | + | 2 |
|  | DMS_other | REaseJ | JACSLS010000010.1_3 | 2401 5391 1 ID=1_3;partial=00;start_type=ATG;rbs_motif=GGAG/GAGG;rbs_spacer=5-10bp;gc_cont=0.419 | JACSLS010000010.1 | 2401 | 5391 | + | 3 |
|  | DMS_other | DrmB | JACSLS010000010.1_4 | 5459 7213 -1 ID=1_4;partial=00;start_type=ATG;rbs_motif=AGGAG;rbs_spacer=5-10bp;gc_cont=0.426 | JACSLS010000010.1 | 5459 | 7213 | - | 4 |
|  | DMS_other | DrmA | JACSLS010000010.1_5 | 7200 10610 -1 ID=1_5;partial=00;start_type=ATG;rbs_motif=GGA/GAG/AGG;rbs_spacer=5-10bp;gc_cont=0.434 | JACSLS010000010.1 | 7200 | 10610 | - | 5 |
|  | argonaute_solo | pAgo | JACSLS010000010.1 12 | 18217 20325 1 ID=1_12;partial=00;start_type=ATG;rbs_motif=GGAG/GAGG;rbs_spacer=5-10bp;gc_cont=0.470 | JACSLS010000010.1 | 18217 | 20325 | + | 12 |
| **sp. MER 193*** | DMS_other | REaseJ | M3591J1280 | M3591_11280 | JAMAW010000021.1 | 750 | 3741 | + | 2 |
|  | DMS_other | DrmB | M3591_11285 | M3591_11285 | JAMAW010000021.1 | 3806 | 5561 | - | 3 |
|  | DMS_other | DrmA | M3591_11290 | M3591_11290 | JAMAVV010000021.1 | 5547 | 8958 | - | 4 |
|  | argonaute_solo | pAgo | M359111320 | M359111320 | JAMAW010000021.1 | 16577 | 18686 | + | 10 |
| **sp. s163*** | argonaute_solo | pAgo | JACSJN010000008.1 _73 | 63997 66102 -1 ID=1_73;partial=00;start_type=ATG;rbs_motif=GGAG/GAGG;rbs_spacer=5-10bp;gc_cont=0.439 | J ACSJ N010000008.1 | 63997 | 66102 | - | 73 |
|  | DMS_other | DrmA | JACSJN010000008.1_79 | 73722 77132 1 ID=1_79;partial=00;start_type=ATG;rbs_motif=GGA/GAG/AGG;rbs_spacer=5-10bp;gc_cont=0.413 | J ACSJ N010000008.1 | 73722 | 77132 | + | 79 |
|  | DMS_other | DrmB | JACSJN010000008.1_80 | 77119 78873 1 ID=1_80;partial=00;start_type=ATG;rbs_motif=AGGAG;rbs_spacer=5-10bp;gc_cont=0.418 | JACSJ N010000008.1 | 77119 | 78873 | + | 80 |
|  | brex_type_l 11 | BrxF | JACSJN010000008.1_81 | 79007 79456 1 ID=1_81;partial=00;start_type=ATG;rbs_motif=GGAGG;rbs_spacer=5-10bp;gc_cont=0.336 | JACSJN010000008.1 | 79007 | 79456 | + | 81 |
|  | brex_type_lll | BrxC | JACSJN010000008.1 _82 | 79478 83182 1 ID=1_82;partial=00;start_type=ATG;rbs_motif=AGGAG;rbs_spacer=5-10bp;gc_cont=0.358 | JACSJ N010000008.1 | 79478 | 83182 | + | 82 |
|  | brex_type_lll | PglXI | JACSJN010000008.1_83 | 83195 85954 1 ID=1_83;partial=00;start_type=ATG;rbs_motif=AGGAG;rbs_spacer=5-10bp;gc_cont=0.329 | JACSJ N010000008.1 | 83195 | 85954 | + | 83 |
|  | brex_type_lll | Pgiz | JACSJN010000008.1_84 | 85973 87859 1 ID=1_84;partial=00;start_type=ATG;rbs_motif=AGGAG/GGAGG;rbs_spacer=11-12bp;gc_cont=0.375 | JACSJ N010000008.1 | 85973 | 87859 | + | 84 |
|  | brex_type_lll | BrxA | JACSJN010000008.1_85 | 87878 88540 1 ID=1_85;partial=00;start_type=ATG;rbs_motif=GGAG/GAGG;rbs_spacer=5-10bp;gc_cont=0.348 | JACSJ N010000008.1 | 87878 | 88540 | + | 85 |
|  | brex_type_lll | BrxHII | JACSJN010000008.1_86 | 88590 91733 1 ID=1_86;partial=00;start_type=ATG;rbs_motif=GGAGG;rbs_spacer=5-10bp;gc_cont=0.331 | JACSJN010000008.1 | 88590 | 91733 | + | 86 |
| **sp. s21*** | wadjet_type_ll | JetA2 | JACSLF010000009.1_31 | 25218 26693 1 ID=1_31;partial=00;start_type=ATG;rbs_motif=AGGAG;rbs_spacer=5-10bp:gc_cont=0.518 | JACSLF010000009.1 | 25218 | 26693 | + | 31 |
|  | wadjet_type_ll | JetB2 | JACSLF010000009.1_32 | 26690 27844 1 ID=1_32;partial=00;start_type=ATG;rbs_motif=GGAG/GAGG;rbs_spacer=5-10bp ;gc_cont=0.514 | JACSLF010000009.1 | 26690 | 27844 | + | 32 |
|  | wadjet_type_ll | JetC2 | JACSLF010000009.1_33 | 27841 31935 1 ID=1_33;partial=00;start_type=ATG;rbs_motif=AGGAG;rbs_spacer=5-10bp;gc_cont=0.538 | JACSLF010000009.1 | 27841 | 31935 | + | 33 |
|  | wadjettypell | JetD2 | JACSLF010000009.1_34 | 31932 33206 1 ID=1_34;partial=00;start_type=ATG;rbs_motif=GGA/GAG/AGG;rbs_spacer=5-10bp;gc_cont=0.538 | JACSLF010000009.1 | 31932 | 33206 | + | 34 |
|  | brex_type_lll | BrxHII | JACSLF010000009.1_36 | 35109 38255 -1 ID=1_36;partial=00;start_type=ATG;rbs_motif=GGAGG;rbs_spacer=5-10bp;gc_cont=0.346 | JACSLF010000009.1 | 35109 | 38255 | - | 36 |
|  | brex_type_lll | BrxA | JACSLF010000009.1_37 | 38305 38967 -1 ID=1_37;partial=00;start_type=ATG;rbs_motif=GGAG/GAGG;rbs_spacer=5-10bp;gc_cont=0.329 | JACSLF010000009.1 | 38305 | 38967 | - | 37 |
|  | brex_type_lll | Pgiz | JACSLF010000009.1_38 | 38986 40875 -1 ID=1_38:partial=00;start_type=ATG:rbs_motif=AGGAG;rbs_spacer=5-10bp:gc_cont=0.379 | JACSLF010000009.1 | 38986 | 40875 | - | 38 |
|  | brextypejll | Pgixi | JACSLF010000009.1_39 | 40891 43656 -1 ID=1_39;partial=00;start_type=ATG;rbs_motif=AGGAG;rbs_spacer=5-10bp;gc_cont=0.332 | JACSLF010000009.1 | 40891 | 43656 | - | 39 |
|  | brextypejll | BrxC | JACSLF010000009.1 _40 | 43669 47373 -1 ID=1_40:partial=00;start_type=ATG;rbs_motif=AGGAG;rbs_spacer=5-10bp;gc_cont=0.361 | JACSLF010000009.1 | 43669 | 47373 | - | 40 |
|  | brextypelll | BrxF | JACSLF010000009.1_41 | 47395 47844 -1 ID=1_41;partial=00;start_type=ATG;rbs_motif=GGAGG;rbs_spacer=5-10bp;gc_cont=0.340 | JACSLF010000009.1 | 47395 | 47844 | - | 41 |
|  | DMS_other | DrmB | JACSLF010000009.1_42 | 47978 49732 -1 ID=1_42;partial=00;start_type=ATG;rbs_motif=AGGAG;rbs_spacer=5-10bp:gc_cont=0.423 | JACSLF010000009.1 | 47978 | 49732 | - | 42 |
|  | DMS_other | DrmA | JACSLF010000009.1_43 | 49719 53129 -1 ID=1_43;partial=00;start_type=ATG;rbs_motif=GGA/GAG/AGG;rbs_spacer=5-10bp;gc_cont=0.431 | JACSLF010000009.1 | 49719 | 53129 | - | 43 |
|  | argonaute_solo | pAgo | JACSLF010000009.1_49 | 60737 62845 1 ID=1_49;partial=00;start_type=ATG;rbs_motif=GGAG/GAGG;rbs_spacer=5-10bp;gc_cont=0.470 | JACSLF010000009.1 | 60737 | 62845 | + | 49 |
| **sp. SH3S1*** | cas_type_l-C | Cas2c | EVJ26_09270 | EVJ26_09270 | SJUW01000007.1 | 42033 | 42324 | - | 39 |
|  | cas_type_l-C | Caslc | EVJ26_09275 | EVJ26_09275 | SJUW01000007.1 | 42333 | 43365 | - | 40 |
|  | cas_type_l-C | Cas4c | EVJ26_09280 | EVJ26_09280 | SJUW01000007.1 | 43361 | 44021 | - | 41 |
|  | cas_type_l-C | Cas7c | EVJ26_09285 | EVJ26_09285 | SJUW01000007.1 | 44010 | 44865 | - | 42 |
|  | cas_type_l-C | Cas8c | EVJ26_09290 | EVJ26_09290 | SJUW01000007.1 | 44861 | 46736 | - | 43 |
|  | cas_type_l-C | Cas5c | EVJ26_09295 | EVJ26_09295 | SJUW01000007.1 | 46748 | 47447 | - | 44 |
|  | cas_type_l-C | Cas3c | EVJ26_09300 | EVJ26_09300 | SJUW01000007.1 | 47462 | 49805 | - | 45 |
|  | argonaute_solo | pAgo | EVJ26_09605 | EVJ26_09605 | SJUW01000007.1 | 98953 | 1E+05 | - | 106 |
|  | brex_type_l | BrxL | EVJ26_09635 | EVJ26_09635 | SJUW01000007.1 | 1E+05 | 1E+05 | - | 112 |
|  | brex_type_l | Pgiz | EVJ26_09640 | EVJ26_09640 | SJUW01000007.1 | 1E+05 | 1E+05 | - | 113 |
|  | brex_type_l | Pgix | EVJ26_09650 | EVJ26_09650 | SJUW01000007.1 | 1E+05 | 1E+05 | - | 115 |
|  | brex_type_l | BrxC | EVJ26_09655 | EVJ26_09655 | SJUW01000007.1 | 1E+05 | 1E+05 | - | 116 |
|  | brex_type_l | BrxB | EVJ26_09660 | EVJ26_09660 | SJUW01000007.1 | 1E+05 | 1E+05 | - | 117 |
|  | brex_type_l | BrxA | EVJ26_09665 | EVJ26_09665 | SJUW01000007.1 | 1E+05 | 1E+05 | - | 118 |
|  | DMS_other | DrmD | EVJ26_09675 | EVJ26_09675 | SJUW01000007.1 | 1E+05 | 1E+05 | - | 120 |
|  | wadjet_type_ll | JetD2 | EVJ26_09690 | EVJ26_09690 | SJUW01000007.1 | 1E+05 | 1E+05 | - | 123 |
|  | wadjet_type_ll | JetC2 | EVJ26_09695 | EVJ26_09695 | SJUW01000007.1 | 1E+05 | 1E+05 | - | 124 |
|  | wadjet_type_ll | JetB2 | EVJ26_09700 | EVJ26_09700 | SJUW01000007.1 | 1E+05 | 1E+05 | - | 125 |
|  | wadjet_type_ll | JetA2 | EVJ26_09705 | EVJ26_09705 | SJUW01000007.1 | 1E+05 | 1E+05 | - | 126 |
| **sp. s55*** | wadjet_type_ll | JetA2 | JACSLE010000011.1_35 | 25058 26533 1 ID=1_35;partial=00;start_type=ATG;rbs_motif=GGAG/GAGG;rbs_spacer=5-10bp;gc_cont=0.522 | JACSLE010000011.1 | 25058 | 26533 | + | 35 |
|  | wadjet_type_ll | JetB2 | JACSLE010000011.1_36 | 26530 27684 1 ID=1_36;partial=00;start_type=ATG;rbs_motif=GGAG/GAGG;rbs_spacer=5-10bp;gc_cont=0.509 | JACSLE010000011.1 | 26530 | 27684 | + | 36 |
|  | wadjet_type_ll | JetC2 | JACSLE010000011.1_37 | 27681 31775 1 ID=1_37;partial=00;start_type=ATG;rbs_motif=AGGAG;rbs_spacer=5-10bp;gc_cont=0.535 | JACSLE010000011.1 | 27681 | 31775 | + | 37 |
|  | wadjet_type_ll | JetD2 | JACSLE010000011.1_38 | 31772 33070 1 ID=1_38;partial=00;start_type=ATG;rbs_motif=GGA/GAG/AGG;rbs_spacer=5-10bp;gc_cont=0.517 | JACSLE010000011.1 | 31772 | 33070 | + | 38 |
|  | brex_type_l | BrxA | JACSLE010000011.1_43 | 37141 37740 1 ID=1_43;partial=00;start_type=ATG;rbs_motif=AGGA;rbs_spacer=5-10bp;gc_cont=0.307 | JACSLE010000011.1 | 37141 | 37740 | + | 43 |
|  | brex_type_l | BrxB | JACSLE010000011.1_44 | 37737 38315 1 ID=1_44:partial=00;start_type=ATG;rbs_motif=AGGAG;rbs_spacer=5-10bp:gc_cont=0.347 | JACSLE010000011.1 | 37737 | 38315 | + | 44 |
|  | brex_type_l | BrxC | JACSLE010000011,1_45 | 38338 41904 1 ID=1_45;partial=00;start_type=ATG;rbs_motif=AGGAG;rbs_spacer=5-10bp;gc_cont=0.385 | JACSLE010000011.1 | 38338 | 41904 | + | 45 |
|  | brex_type_l | PglX | JACSLE010000011.1_46 | 41916 45437 1 ID=1_46:partial=00;start_type=ATG;rbs_motif=GGA/GAG/AGG;rbs_spacer=5-10bp;gc_cont=0.338 | JACSLE010000011.1 | 41916 | 45437 | + | 46 |
|  | brex_type_l | Pgiz | JACSLE010000011.1_48 | 46378 48936 1 ID=1_48;partial=00;start_type=GTG;rbs_motif=GGAG/GAGG;rbs_spacer=5-10bp;gc_cont=0.369 | JACSLE010000011.1 | 46378 | 48936 | + | 48 |
|  | brex_type_l | BrxL | JACSLE010000011.1_49 | 48955 51000 1 ID=1_49;partial=00;start_type=ATG;rbs_motif=AGGAG;rbs_spacer=5-10bp;gc_cont=0.448 | JACSLE010000011.1 | 48955 | 51000 | + | 49 |
|  | argonaute_solo | pAgo | JACSLE010000011.1_55 | 60030 62138 1 ID=1_55;partial=00;start_type=ATG;rbs_motif=GGAG/GAGG;rbs_spacer=5-10bp;gc_cont=0.471 | JACSLE010000011.1 | 60030 | 62138 | + | 55 |
|  | cas_type_ll-A | Cas9 | JACSLE010000011.1_93 | 87360 91523 1 ID=1_93;partial=00;start_type=ATG;rbs_motif=AGGAG;rbs_spacer=5-10bp;gc_cont=0.371 | JACSLE010000011.1 | 87360 | 91523 | + | 93 |
|  | cas_type_ll-A | Cas1_ll | JACSLE010000011.1_94 | 91539 92420 1 ID=1_94;partial=00;start_type=ATG;rbs_motif=GGA/GAG/AGG;rbs_spacer=5-10bp;gc_cont=0.374 | JACSLE010000011.1 | 91539 | 92420 | + | 94 |
|  | cas_type_ll-A | Cas2_ll | JACSLE010000011.1_95 | 92417 92737 1 ID=1_95;partial=00;start_type=ATG;rbs_motif=AGGAG;rbs_spacer=5-10bp;gc_cont=0.364 | JACSLE010000011.1 | 92417 | 92737 | + | 95 |
|  | cas_type_ll-A | Csn2 | JACSLE010000011.1_96 | 92734 93603 1 ID=1_96;partial=00;start_type=ATG;rbs_motif=GGA/GAG/AGG;rbs_spacer=5-10bp;gc_cont=0.351 | JACSLE010000011.1 | 92734 | 93603 | + | 96 |
| **sp. CinTr1*** | argonautesolo | pAgo | CAN DNX010000009.1_71 | 62648 64756 -1 ID=1_71:partial=00;start_type=ATG;rbs_motif=GGAG/GAGG;rbs_spacer=5-10bp;gc_cont=0.450 | CAN DNX010000009.1 | 62648 | 64756 | - | 71 |
|  | RM_type_l | REase_l | CAN DNX010000009.1_77 | 78296 81286 -1 ID=1_77;partial=00;start_type=ATG;rbs_motif=GGAG/GAGG;rbs_spacer=5-10bp;gc_cont=0.417 | CAN DNX010000009.1 | 78296 | 81286 | - | 77 |
|  | RM_type_l | Specificity_l | CAN DNX010000009.1_78 | 81301 82635 -1 ID=1_78;partial=00;start_type=ATG;rbs_motif=AGxAGG/AGGxGG;rbs_spacer=5-10bp;gc_cont=0.321 | CAN DNX010000009.1 | 81301 | 82635 | - | 78 |
|  | RM_type_l | MTaseJ | CAN DNX010000009.1_79 | 82632 84644 -1 ID=1_79;partial=00;start_type=ATG;rbs_motif=GGAG/GAGG;rbs_spacer=5-10bp;gc_cont=0.382 | CAN DNX010000009.1 | 82632 | 84644 | - | 79 |
|  | LamassuFamily | LmuA | CAN DNX010000009.1_114 | 112876114072 1 ID=1_114:partial=00:start_type=ATG;rbs_motif=AGGA/GGAG/GAGG;rbs_spacer=11- 12bp;gc_cont=0.317 | CANDNX010000009.1 | 1E+05 | 1E+05 | + | 114 |
|  | LamassuFamily | LmuC | CAN DNX010000009.1_115 | 114069 114560 1 ID=1_115;partial=00;start_type=ATG;rbs_motif=GGA/GAG/AGG;rbs_spacer=5-10bp;gc_cont=0.321 | CAN DNX010000009.1 | 1E+05 | 1E+05 | + | 115 |
|  | Lamassu_Family | LmuB | CAN DNX010000009.1_116 | 114561 116486 1 ID=1_116;partial=00;start_type=GTG;rbs_motif=AGGA/GGAG/GAGG;rbs_spacer=11-12bp;gc_cont=0.338 | CANDNX010000009.1 | 1E+05 | 1E+05 | + | 116 |
|  | Lamassu_Family | LmuB | CAN DNX010000009.1_117 | 116723 117697 -1 ID=1_117;partial=00;start_type=ATG:rbs_motif=AGGAG;rbs_spacer=5-10bp;gc_cont=0.456 | CANDNX010000009.1 | 1E+05 | 1E+05 | - | 117 |
| **sp. s7**** | argonaute_solo | pAgo | JACSLO010000010.1_71 | # 55157 # 57265 # -1 # ID=1_71 ;partial=00;start_type=ATG;rbs_motif=GGAG/GAGG;rbs_spacer=5-10bp;gc_cont=0.479 | JACSLO010000010.1 | 55157 | 57265 | - | 71 |
|  | PDC-S18 | PDC-S18 | JACSL0010000010.1_72 | # 57414 # 58313 # -1 # ID=1_72;partial=00;start_type=ATG;rbs_motif=AGGAG;rbs_spacer=5-10bp;gc_cont=0.420 | JACSLO010000010.1 | 57414 | 58313 | - | 72 |
|  | RMJypeJ | REaseJ | JACSLO010000010.1_80 | # 75504 # 78494 # -1 # ID=1_80;partial=00;start_type=ATG;rbs_motif=GGAG/GAGG;rbs_spacer=5-10bp;gc_cont=0.415 | JACSLO010000010.1 | 75504 | 78494 | - | 80 |
|  | RM_type_l | SpecificityJ | JACSLO010000010.1_81 | # 78510 # 79751 # -1 # ID=1_81 ;partial=00;start_type=ATG;rbs_motif=AGxAGG/AGGxGG;rbs_spacer=5-10bp;gc_cont=0.324 | JACSLO010000010.1 | 78510 | 79751 | - | 81 |
|  | RM_type_l | MTase_l | JACSLO010000010.1_82 | # 79748 # 81760 # -1 # ID=1_82;partial=00;start_type=ATG;rbs_motif=GGAG/GAGG;rbs_spacer=5-10bp;gc_cont=0.380 | J ACSLO010000010.1 | 79748 | 81760 | - | 82 |
| **sp. AT1b**** | argonaute_solo | pAgo | MOEL01000004.1_65 | # 53075 # 55183 # -1 # ID=1_65;partial=00;start_type=ATG;rbs_motif=GGAG/GAGG;rbs_spacer=5-10bp;gc_cont=0.446 | MOEL01000004.1 | 53075 | 55183 | - | 65 |
|  | RM_type_l | REase_l | MOEL01000004.1 _71 | # 68724 # 71714 # -1 # ID=1_71 ;partial=00;start_type=ATG;rbs_motif=GGAG/GAGG;rbs_spacer=5-10bp;gc_cont=0.417 | MOEL01000004.1 | 68724 | 71714 | - | 71 |
|  | RM_type_l | Specificity_l | MOEL01000004.1_72 | # 71729 # 73027 # -1 # ID=1_72;partial=00;start_type=GTG;rbs_motif=AGxAGG/AGGxGG;rbs_spacer=5-10bp;gc_cont=0.324 | MOEL01000004.1 | 71729 | 73027 | - | 72 |
|  | RM_type_l | MTase_l | MOEL01000004.1_73 | # 73024 # 75036 # -1 # ID=1_73;partial=00;start_type=ATG;rbs_motif=None;rbs_spacer=None;gc__cont=0.378 | MOEL01000004.1 | 73024 | 75036 | - | 73 |
| **sp. JLM-2**** | argonaute_solo | pAgo | LDNY01000004.1_87 | 78280 80058 -1 ID=1_87;partial=00;start_type=ATG;rbs_motif=None;rbs_spacer=none;gc_cont=0.449 | LDNY01000004.1 | 78280 | 80058 | - | 87 |
| **sp. N5**** | argonaute_solo | pAgo | OKS35_14005 | - | JAPAEP010000004.1 | 78744 | 80853 | - | 91 |
|  | PD-T4-6 | PD-T4-6 | OKS35_14020 | - | JAPAEP010000004.1 | 83115 | 83937 | - | 94 |
| **sp. s48**** | argonaute_solo | pAgo | J ACSKC010000004.1 _90 | # 79247 # 81352 # -1 # ID=1_90;partial=00;start_type=ATG;rbs_motif=GGAG/GAGG;rbs_spacer=5-10bp;gc_cont=0.443 | JACSKC010000004.1 | 79247 | 81352 | - | 90 |
|  | hachiman_typej | HamA1 | JACSKC010000004.1 _94 | # 82897 # 83724 # 1 # ID=1_94;partial=00;start_type=ATG;rbs_motif=AGGA;rbs_spacer=5-10bp;gc_cont=0.345 | JACSKC010000004.1 | 82897 | 83724 | + | 94 |
|  | hachiman_type_l | HamB1 | JACSKC010000004.1 _95 | # 83714 # 86833 # 1 # ID=1_95;partial=00;start_type=ATG;rbs_motif=AGGAG;rbs_spacer=5-10bp;gc_cont=0.324 | JACSKC010000004.1 | 83714 | 86833 | + | 95 |
| ***aestuarii******* | argonaute_solo | pAgo | LCM29_13835 | - | JAIVAC010000004.1 | 36145 | 38251 | + | 40 |
|  | cas_type_l-C | Cas3c | LCM29_14100 | - | JAIVAC010000004.1 | 75868 | 78202 | + | 93 |
|  | cas_type_l-C | Cas5c | LCM29_14105 | - | JAIVAC010000004.1 | 78218 | 78917 | + | 94 |
|  | cas_type_l-C | Cas8c | LCM29_14110 | - | JAIVAC010000004.1 | 78928 | 80806 | + | 95 |
|  | cas_type_l-C | Cas7c | LCM29_14115 | - | JAIVAC010000004.1 | 80802 | 81657 | + | 96 |
|  | cas_type_l-C | Cas4c | LCM29_14120 | - | JAIVAC010000004.1 | 81646 | 82306 | + | 97 |
|  | cas_type_l-C | Caslc | LCM29_14125 | - | JAIVAC010000004.1 | 82302 | 83334 | + | 98 |
|  | cas_type_l-C | Cas2c | LCM29_14130 | - | JAIVAC010000004.1 | 83344 | 83635 | + | 99 |
| ***marinum******* | Mokosh_TypeII | MkoC | JHZT01000001.1 _40 | # 38434 # 41529 # -1 # ID=1_40;partial=00;start_type=GTG;rbs_motif=AGGAG;rbs_spacer=5-10bp;gc_cont=0.372 | JHZT01000001.1 | 38434 | 41529 | - | 40 |
|  | argonaute_solo | pAgo | JHZT01000001.1_98 | # 92041 # 94146 # -1 # ID=1_98;partial=00;start_type=ATG;rbs_motif=AGGA;rbs_spacer=5-10bp;gc_cont=0.445 | JHZT01000001.1 | 92041 | 94146 | - | 98 |
|  | GAO_19 | HerA | JHZT01000001.1_101 | # 95465 # 96595 # -1 # ID=1_101;partial=00;start_type=ATG;rbs_motif=None;rbs_spacer=None;gc_cont=0.354 | JHZT01000001.1 | 95465 | 96595 | - | 101 |
|  | GAO_19 | HerA | J HZT01000001.1_102 | # 96622 # 97209 # -1 # ID=1_102:partial=00;start_type=ATG;rbs_motif=AGGA/GGAG/GAGG;rbs_spacer=11-12bp;gc_cont=0.320 | JHZT01000001.1 | 96622 | 97209 | - | 102 |
|  | GAO_19 | SIR2 | JHZT01000001.1_103 | # 97210 # 98277 # -1 # ID=1_103;partial=00;start_type=ATG;rbs_motif=AGxAGG/AGGxGG;rbs_spacer=5-10bp;gc_cont=0.316 | JHZT01000001.1 | 97210 | 98277 | - | 103 |
|  | RM_type_ll | MTaseJI | JHZT01000001,1_370 | # 351406 # 352740 # -1 # ID=1_370;partial=00;start_type=TTG;rbs_motif=GGA/GAG/AGG;rbs_spacer=5-10bp;gc_cont=0.495 | JHZT01000001.1 | 4E+05 | 4E+05 | - | 370 |
|  | VSPR | MTase_ll | J HZT01000001.1_370 | # 351406 # 352740 # -1 # ID=1_370;partial=00;start_type=TTG;rbs_motif=GGA/GAG/AGG;rbs_spacer=5-10bp;gc_cont=0.495 | JHZT01000001.1 | 4E+05 | 4E+05 | - | 370 |
|  | RM_type_ll | REaseJI | JHZT01000001.1_371 | # 352900 # 354270 # 1 # ID=1_371;partial=00;start_type=TTG;rbs_motif=AGGAG;rbs_spacer=5-10bp;gc_cont=0.448 | JHZT01000001.1 | 4E+05 | 4E+05 | + | 371 |
|  | VSPR | Vsr | JHZT01000001.1_372 | # 354326 # 354772 # 1 # ID=1_372;partial=00;start_type=ATG;rbs_motif=GGAGG;rbs_spacer=5-10bp;gc_cont=0.479 | JHZT01000001.1 | 4E+05 | 4E+05 | + | 372 |
|  | PDC-S07 | PDC-S07 | J HZT01000001.1 _385 | # 365842 # 366255 # 1 # ID=1_385;partial=00;start_type=ATG;rbs_motif=GGAG/GAGG;rbs_spacer=5-10bp;gc_cont=0.362 | JHZT01000001.1 | 4E+05 | 4E+05 | + | 385 |
|  | RM_type_IV | mREaseJV | JHZT01000001.1_386 | # 366275 # 369154 # -1 # ID=1_386;partial=00;start_type=ATG;rbs_motif=AGGAG;rbs_spacer=5-10bp;gc_cont=0.337 | JHZT01000001.1 | 4E+05 | 4E+05 | - | 386 |
| **sp. s38**** | argonaute_solo | pAgo | JACSKT010000005.1_90 | 83375 84274 -1 ID=1_90;partiaI=00;start_type=ATG;rbs_motif=GGA/GAG/AGG;rbs_spacer=5-10bp;gc_cont=0.473 | JACSKT010000005.1 | 83375 | 84274 | - | 90 |
|  | argonaute_solo | pAgo | JACSKT010000005.1_91 | 84234 85484 -1 ID=1_91;partial=00;start_type=ATG;rbs_motif=GGAG/GAGG;rbs_spacer=5-10bp;gc_cont=0.460 | JACSKT010000005.1 | 84234 | 85484 | - | 91 |
|  | Lamassu_Family | LmuB | JACSKT010000005.1_93 | 87264 90410 -1 ID=1_93;partial=00;start_type=ATG;rbs_motif=GGAG/GAGG;rbs_spacer=5-10bp;gc_cont=0.262 | JACSKT010000005.1 | 87264 | 90410 | - | 93 |
|  | Lamassu_Family | LmuC | JACSKT010000005.1_94 | 90403 90987 -1 ID=1_94;partiaI=00;start_type=TTG;rbs_motif=AGGAG;rbs_spacer=5-10bp;gc_cont=0.246 | JACSKT010000005.1 | 90403 | 90987 | - | 94 |
|  | Lamassu_Family | LmuA | JACSKT010000005.1_95 | 91002 92336 -1 ID=1_95;partial=00;start_type=ATG;rbs_motif=GGAG/GAGG;rbs_spacer=5-10bp;gc_cont=0.285 | JACSKT010000005.1 | 91002 | 92336 | - | 95 |
|  | Lamassu_Family | LmuA | JACSKT010000005.1_96 | 92315 93406 -1 ID=1_96;partiaI=00;start_type=GTG;rbs_motif=GGAGG;rbs_spacer=5-10bp;gc_cont=0.297 | JACSKT010000005.1 | 92315 | 93406 | - | 96 |
| **sp. s89**** | argonaute_solo | pAgo | JACSKS010000005.1_90 | # 83375 # 84274 # -1 # ID=1_90;partial=00;start_type=ATG;rbs_motif=GGA/GAG/AGG;rbs_spacer=5-10bp;gc_cont=0.473 | JACSKS010000005.1 | 83375 | 84274 | - | 90 |
|  | argonaute_solo | pAgo | JACSKS010000005.1_91 | # 84234 # 85484 # -1 # ID=1_91 ;partial=00;start_type=ATG;rbs_motif=GGAG/GAGG;rbs_spacer=5-10bp;gc_cont=0.460 | JACSKS010000005.1 | 84234 | 85484 | - | 91 |
|  | Lamassu_Family | LmuB | JACSKS010000005.1_93 | # 87264 # 90410 # -1 # ID=1_93;partial=00;start_type=ATG;rbs_motif=GGAG/GAGG;rbs_spacer=5-10bp;gc_cont=0.262 | JACSKS010000005.1 | 87264 | 90410 | - | 93 |
|  | Lamassu_Family | LmuC | JACSKS010000005.1_94 | # 90403 # 90987 # -1 # ID=1_94;partial=00;start_type=TTG;rbs_motif=AGGAG;rbs_spacer=5-10bp;gc_cont=0.246 | JACSKS010000005.1 | 90403 | 90987 | - | 94 |
|  | Lamassu_Family | LmuA | JACSKS010000005.1_95 | # 91002 # 92336 # -1 # ID=1_95;partial=00;start_type=ATG;rbs_motif=GGAG/GAGG;rbs_spacer=5-10bp;gc_cont=0.285 | JACSKS010000005.1 | 91002 | 92336 | - | 95 |
|  | Lamassu_Family | LmuA | JACSKS010000005.1_96 | # 92315 # 93406 # -1 # ID=1_96;partial=00;start_type=GTG;rbs_motif=GGAGG;rbs_spacer=5-10bp;gc_cont=0.297 | JACSKS010000005.1 | 92315 | 93406 | - | 96 |
| **sp. s139**** | Lamassu_Family | LmuA | JACSKU010000005.1 _30 | # 21736 # 22827 # 1 # ID=1_30;partial=00;start_type=GTG;rbs_motif=GGAGG;rbs_spacer=5-10bp;gc_cont=0.297 | JACSKU010000005.1 | 21736 | 22827 | + | 30 |
|  | Lamassu_Family | LmuA | JACSKU010000005.1_31 | # 22806 # 24140 # 1 # ID=1_31 ;partial=00;start_type=ATG;rbs_motif=GGAG/GAGG;rbs_spacer=5-10bp;gc_cont=0.286 | JACSKU010000005.1 | 22806 | 24140 | + | 31 |
|  | Lamassu_Family | LmuC | JACSKU010000005.1_32 | # 24155 # 24739 # 1 # ID=1_32;partial=00;start_type=TTG;rbs_motif=AGGAG;rbs_spacer=5-10bp;gc_cont=0.246 | JACSKU010000005.1 | 24155 | 24739 | + | 32 |
|  | Lamassu_Family | LmuB | JACSKU010000005.1_33 | # 24732 # 27878 # 1 # ID=1_33;partial=00;start_type=ATG;rbs_motif=GGAG/GAGG;rbs_spacer=5-10bp;gc_cont=0.262 | JACSKU010000005.1 | 24732 | 27878 | + | 33 |
|  | argonaute_solo | pAgo | JACSKU010000005.1_35 | # 29658 # 30908 # 1 # ID=1_35;partial=00;start_type=ATG;rbs_motif=GGAG/GAGG;rbs_spacer=5-10bp;gc_cont=0.460 | JACSKU010000005.1 | 29658 | 30908 | + | 35 |
|  | argonaute_solo | pAgo | JACSKU010000005.1_36 | # 30868 # 31767 # 1 # ID=1_36;partial=00;start_type=ATG;rbs_motif=GGA/GAG/AGG;rbs_spacer=5-10bp;gc_cont=0.471 | JACSKU010000005.1 | 30868 | 31767 | + | 36 |
| **sp. s196**** | argonaute_solo | pAgo | JACSJR010000010.1_42 | 33289 35397 1 ID=1_42;partial=00;start_type=ATG;rbs_motif=AGGA;rbs_spacer=5-10bp;gc_cont=0.480 | JACSJR010000010.1 | 33289 | 35397 | + | 42 |
|  | cas_type_l-C | Cas3c | JACSJR010000010.1_79 | 62756 65089 1 ID=1_79;partial=00;start_type=GTG;rbs_motif=GGA/GAG/AGG;rbs_spacer=5-10bp;gc_cont=0.392 | JACSJR010000010.1 | 62756 | 65089 | + | 79 |
|  | cas_type_l-C | Cas5c | JACSJR010000010.1_80 | 65106 65804 1 ID=1_80;partial=00;start_type=ATG;rbs_motif=AGGAG;rbs_spacer=5-10bp;gc_cont=0.391 | JACSJR010000010.1 | 65106 | 65804 | + | 80 |
|  | cas_type_l-C | Cas8c | JACSJR010000010.1_81 | 65816 67693 1 ID=1_81;partial=00;start_type=ATG;rbs_motif=AGGAG;rbs_spacer=5-10bp;gc_cont=0.412 | JACSJR010000010.1 | 65816 | 67693 | + | 81 |
|  | cas_type_l-C | Cas7c | JACSJR010000010.1_82 | 67690 68544 1 ID=1_82;partial=00;start_type=ATG;rbs_motif=AGGAG/GGAGG;rbs_spacer=11-12bp;gc_cont=0.416 | JACSJR010000010.1 | 67690 | 68544 | + | 82 |
|  | cas_type_l-C | Cas4c | JACSJR010000010.1_83 | 68534 69193 1 ID=1_83;partial=00;start_type=ATG;rbs_motif=None;rbs_spacer=None;gc_cont=0.409 | JACSJR010000010.1 | 68534 | 69193 | + | 83 |
|  | cas_type_l-C | Caslc | JACSJR010000010.1_84 | 69190 70221 1 ID=1_84;partial=00;start_type=TTG;rbs_motif=AGGAG;rbs_spacer=5-10bp;gc_cont=0.396 | JACSJR010000010.1 | 69190 | 70221 | + | 84 |
|  | cas_type_l-C | Cas2c | JACSJR010000010.1_85 | 70232 70522 1 ID=1_85;partial=00;start_type=ATG;rbs_motif=GGAG/GAGG;rbs_spacer=5-10bp;gc_cont=0.388 | JACSJR010000010.1 | 70232 | 70522 | + | 85 |
| **sp. s149**** | argonaute_solo | pAgo | JACSLZ010000009.1_32 | # 24185 # 26293 # 1 # ID=1_32;partial=00;start_type=ATG;rbs_motif=GGAG/GAGG;rbs_spacer=5-10bp;gc_cont=0.467 | JACSLZ010000009.1 | 24185 | 26293 | + | 32 |
|  | cas_type_l-C | Cas3c | JACSLZ010000009.1_79 | # 54284 # 56632 # 1 # ID=1_79;partial=00;start_type=GTG;rbs_motif=GGAG/GAGG;rbs_spacer=5-10bp;gc_cont=0.413 | JACSLZ010000009.1 | 54284 | 56632 | + | 79 |
|  | cas_type_l-C | Cas5c | JACSLZ010000009.1_80 | # 56648 # 56983 # 1 # ID=1_80;partial=00;start_type=ATG;rbs_motif=AGGAG;rbs_spacer=5-10bp;gc_cont=0.438 | JACSLZ010000009.1 | 56648 | 56983 | + | 80 |
|  | cas_type_l-C | Cas8c | JACSLZ010000009.1_81 | # 57358 # 59232 # 1 # ID=1_81 ;partial=00;start_type=ATG;rbs_motif=AGGAG;rbs_spacer=5-10bp;gc_cont=0.421 | JACSLZ010000009.1 | 57358 | 59232 | + | 81 |
|  | cas_type_l-C | Cas7c | JACSLZ010000009.1_82 | # 59229 # 60083 # 1 # ID=1_82;partial=00;start_type=ATG;rbs_motif=GGA/GAG/AGG;rbs_spacer=5-10bp;gc_cont=0.440 | JACSLZ010000009.1 | 59229 | 60083 | + | 82 |
|  | cas_type_l-C | Cas4c | JACSLZ010000009.1_83 | # 60073 # 60732 # 1 # ID=1_83;partial=00;start_type=ATG;rbs_motif=None;rbs_spacer=None;gc_cont=0.418 | JACSLZ010000009.1 | 60073 | 60732 | + | 83 |
|  | cas_type_l-C | Caslc | JACSLZ010000009.1_84 | # 60738 # 61760 # 1 # ID=1_84;partial=00;start_type=ATG;rbs_motif=GGA/GAG/AGG;rbs_spacer=5-10bp;gc_cont=0.432 | JACSLZ010000009.1 | 60738 | 61760 | + | 84 |
|  | cas_type_l-C | Cas2c | JACSLZ010000009.1_85 | # 61770 # 62060 # 1 # ID=1_85;partial=00;start_type=ATG;rbs_motif=GGAG/GAGG;rbs_spacer=5-10bp;gc_cont=0.395 | JACSLZ010000009.1 | 61770 | 62060 | + | 85 |
| **sp. SL-10**** | wadjet_type_ll | JetA2 | EVJ33_10150 | - | SJUP01000008.1 | 36463 | 37939 | + | 41 |
|  | wadjet_type_ll | JetB2 | EVJ33_10155 | - | SJUP01000008.1 | 37935 | 39090 | + | 42 |
|  | wadjet_type_ll | JetC2 | EVJ33_10160 | - | SJUP01000008.1 | 38978 | 43181 | + | 43 |
|  | wadjet_type_ll | JetD2 | EVJ33_10165 | - | SJUP01000008.1 | 43135 | 44440 | + | 44 |
|  | RM_type_l | MTase_l | EVJ33_10175 | - | SJUP01000008.1 | 45836 | 47423 | + | 46 |
|  | RM_type_l | Specificity_l | EVJ33_10180 | - | SJUP01000008.1 | 47419 | 48568 | + | 47 |
|  | RM_type_l | REase_l | EVJ33_10185 | - | SJUP01000008.1 | 48592 | 51760 | + | 48 |
|  | argonaute_solo | pAgo | EVJ33_10200 | - | SJUP01000008.1 | 56821 | 58930 | + | 51 |
| **sp. s181**** | argonaute_solo | pAgo | JACSJ M010000005.1 _72 | 63916 66024 -1 ID-1_72;partial=00;start_type=ATG;rbs_rnotif-AGGA;rbs_spacer-5-10bp;gc_cx;nt-0.480 | JACSJM010000005.1 | 63916 | 66024 | - | 72 |
| **sp. s140**** | argonaute_solo | pAgo | JACSJK010000006.1_75 | # 64567 # 66672 # -1 # ID=1_75;partial=00;start_type=ATG;rbs_motif=GGAG/GAGG;rbs_spacer=5-10bp;gc_cont=0.440 | JACSJK010000006.1 | 64567 | 66672 | - | 75 |
|  | PDC-S18 | PDC-S18 | JACSJK010000006.1 _76 | # 66824 # 67723 # -1 # ID=1_76;partial=00;start_type=ATG;rbs_motif=AGGAG;rbs_spacer=5-10bp;gc_cont=0.416 | JACSJK010000006.1 | 66824 | 67723 | - | 76 |
|  | RM_type_l | MTase_l | JACSJK010000006.1_78 | # 70152 # 72035 # 1 # ID=1_78;partial=00;start_type=ATG;rbs_motif=AGGAG;rbs_spacer=5-10bp;gc_cont=0.340 | JACSJK010000006.1 | 70152 | 72035 | + | 78 |
|  | RM_type_l | MTaseJ | JACSJK010000006.1_79 | # 72473 # 73993 # 1 # ID=1_79;partial=00:start_type=ATG;rbs_motif=AGxAGG/AGGxGG;rbs_spacer=5-10bp;gc_cont=0.408 | JACSJK010000006.1 | 72473 | 73993 | + | 79 |
|  | RMJypeJ | Specificity_l | JACSJK010000006.1_80 | # 73983 # 75230 # 1 # ID=1_80;partial=00;start_type=ATG;rbs_motif=GGA/GAG/AGG;rbs_spacer=11-12bp;gc_cont=0.321 | JACSJK010000006.1 | 73983 | 75230 | + | 80 |
|  | RM_type_l | REaseJ | JACSJK010000006.1_81 | # 75234 # 78344 # 1 # ID=1_81;partial=00;start_type=ATG;rbs_motif=GGAGG;rbs_spacer=5-10bp;gc_cont=0.386 | JACSJK010000006.1 | 75234 | 78344 | + | 81 |
|  | argonaute_solo | pAgo | QLVE01000025.1_7 | # 10539 # 12647 # 1 # ID=1_7;partial=00;start_type=ATG;rbs_motif=GGAG/GAGG;rbs_spacer=5-10bp;gc_cont=0.472 | QLVE01000025.1 | 10539 | 12647 | + | 7 |
|  | PDC-S18 | PDC-S18 | JACSLJ010000015.1_4 | #2671 # 3570 # -1 # ID=1_4;partial=00;start_type=ATG;rbs_motif=GGAG/GAGG;rbs_spacer=5-10bp;gc_cont=0.413 | JACSLJ010000015.1 | 2671 | 3570 | - | 4 |

“*” Indicates contigs annotated in September of 2023. “**” Indicates contigs annotated in April of 2024. Several uncharacterized candidate defense systems (e.g. PDC systems) were identified by PADLOC on these more recently annotated contigs; however, these were not included for any subsequent analysis.

**Supplementary Table 3. Experimentally characterized pAgos derived from various bacterial genera used for bioinformatics analyses in this study.**

| **Host organism** | **pAgo name** | **pAgo NCBI accession** | **Source** |
| --- | --- | --- | --- |
| *Aquifex aeolicus* | AaAgo | WP_010880937.1 | [64] |
| *Archaeoglobus fulgidus* | AfAgo | WP_010878815.1 | [65] |
| *Aromatoleum aromaticum* EbN1 | ArAgo | CAI09102.1 | [38] |
| *Butyrivibrio* sp. INlla16 | BsAgo | WP_091870084.1 |  |
| *Clostridium saudiense* | CaAgo | WP_042399050.1 |  |
| *Clostridium butyricum* | CbAgo | WP_058142162.1 | [17] |
| *Clostridium perfringens* WAL-14572 | CpAgo | EHP50500.1 | [39] |
| *Clostridium sartagoforme* | CsAgo | WP_016205751.1 | [38] |
| *Dorea longicatena* | DlAgo | WP_055195547.1 | [18] |
| *Fusicatenibacter saccharivorans* | FsAgo | WP_055268308.1 | [38] |
| *Fischerella thermalis* | FtAgo | WP_009456812.1 |  |
| *Intestinibacter bartlettii* DSM 16795 | IbAgo | WP_055087491.1 | [39] |
| *Kurthia massiliensis* | KmAgo | WP_010289662.1 | [40] |
| *Limnothrix rosea* | LrAgo | WP_075892274.1 | [24] |
| *Lyngbya* sp. PCC 8106 | LsAgo | WP_009787621.1 | [38] |
| *Microcystis aeruginosa* | MaAgo | WP_061432364.1 |  |
| *Methylotuvimicrobium buryatense* | MbAgo | WP_017841076.1 |  |
| *Methanocaldococcus jannaschii* | MjAgo | WP_010870838.1 | [16] |
| *Marinitoga piezophila* | MpAgo | WP_014295921.1 | [66] |
| *Pyrococcus furiosus* | PfAgo | WP_011011654.1 | [14] |
| *Prochlorothrix hollandica* PCC 9006 = CALU 1027 | PhAgo | KKI99518.1 | [38] |
| *Pseudoalteromonas luteoviolacea* B = ATCC 29581 | PuAgo | CCQ09996.1 |  |
| *Pseudobutyrivibrio xylanivorans* | PxAgo | WP_090164194.1 |  |
| *Rhodobacter sphaeroides* | RsAgo | WP_011910606.1 | [15] |
| *Thermus thermophilus* | TtAgo | WP_014511275.1 | [13] |

**Supplementary Table 4. Guide strands used in this study.**

| **Name** | **Sequence (5’-3’)** | **Description** | **Remarks** |
| --- | --- | --- | --- |
| **pQE-80L guides** | | |  |
| SML657 | /5Phos/TAACTACGGCTACACTAGAAG | F, targets pQE-80L *Ori*. |  |
| SML658 | /5Phos/TGTCCTTCTAGTGTAGCCGTA | R, targets pQE-80L *Ori*. Produces sticky ends with SML657. |  |
| SML659 | /5Phos/TCATGGGAGAAAATAATACTG | F, targets pQE-80L *lacI*. | 5’-OH version of this guide was also used |
| SML660 | /5Phos/TCAACAGTATTATTTTCTCCC | R, targets pQE-80L *lacI*. Produces sticky ends with SML659. | 5’-OH version of this guide was also used |
| **M13 genome guides** | | |  |
| SML708 | /5Phos/TATCAACAATAGATAAGTCCT | Targets M13 site 1 (~3.6k bases). |  |
| SML709 | /5Phos/TTCCTCGTTAGAATCAGAGCG | Targets M13 site 2 (~5.4k bases). |  |
| **Other guides** | | |  |
| SML661 | /5Phos/GCCAGCTAATCGACTCGTCGA | “Scrambled”/NTC guide. |  |

Guides are 21 nt-long ssDNA oligos with 5’ phosphate groups and 5’-terminal thymine nucleotides. 5’-hydroxylated versions of SML659 and SML660 were also used.
